# Cortical evidence accumulation for perceptual experience occurs irrespective of reports

**DOI:** 10.1101/2024.03.20.585198

**Authors:** François Stockart, Ramla Msheik, Alexis Robin, Lenka Jurkovičová, Dorian Goueytes, Martin Rouy, Radek Mareček, Dominique Hoffmann, Liad Mudrik, Robert Roman, Milan Brázdil, Lorella Minotti, Philippe Kahane, Michael Pereira, Nathan Faivre

**Affiliations:** Univ. Grenoble Alpes, Univ. Savoie Mont Blanc, CNRS, LPNC, 38000 Grenoble, France; Neurology Department, CHU Grenoble Alpes, Univ. Grenoble Alpes, Inserm, U121, Grenoble Institut Neurosciences, 3800 Grenoble, France; 1st Department of Neurology, St. Anne Univ. Hospital and Faculty of Medicine, Masaryk University, Brno, Czech Republic; Behavioral and Social Neuroscience Research Group, CEITEC MU, Brno, Czech Republic; Neurosurgery Department, CHU Grenoble Alpes, Univ. Grenoble Alpes, Inserm, U121, Grenoble Institut Neurosciences, 3800 Grenoble, France; School of Psychological Sciences, Tel Aviv University, Tel Aviv, Israel; Sagol School of Neuroscience, Tel Aviv University, Tel Aviv, Israel; Univ. Grenoble Alpes, Inserm, Grenoble Institut Neurosciences, 38000 Grenoble, France

**Keywords:** intracranial EEG, evidence accumulation, perceptual experience, conscious access, perceptual monitoring, confidence, no report, ventral visual cortex, multivariate decoding, computational modeling

## Abstract

Perceptual experience is a multi-faceted, dynamical process, tackled empirically through measures of stimulus detectability and confidence. To assess if stimulus detection and confidence can be explained by evidence accumulation, a form of sequential sampling of sensory evidence, we analyzed high-gamma activity from stereo-electroencephalographic data of 29 participants partaking in 3 pre-registered experiments. In an immediate-response experiment, individual channels and decoded multivariate latent variables in the visual, inferior frontal, and anterior insular cortices displayed functional markers of evidence accumulation. In two further experiments, this signal in the ventral visual cortex differentiated between (1) seen and unseen stimuli in delayed detection, (2) high and low intensity stimuli during passive viewing, and (3) levels of confidence when stimuli were seen. A computational model of leaky evidence accumulation successfully reproduced both behavioral and neural data. Overall, these results indicate that evidence accumulation explains key aspects of perceptual experience, encompassing both conscious access and monitoring.

## Introduction

Perceptual experience is a dynamic phenomenon, with percepts continuously coming in and out of our stream of consciousness (James, 1890). In addition, monitoring these percepts leads to varying levels of confidence, which is held by some to be an integral part of perceptual experience (Dokic & Martin, 2015; Pereira et al., 2022). What processes in the brain determine when a percept is consciously accessed, and monitor the quality of sensory evidence underlying that percept? The most common approach to studying perceptual experience does not address these questions, as it only purports to explain stimulus detection, i.e., what happens in the brain if a stimulus has already reached conscious access (Crick & Koch, 1990; Koch et al., 2016). Instead, one proposed mechanism to explain the temporal dynamics leading to a perceptual experience is evidence accumulation (Pereira et al., 2022). In evidence accumulation models, the brain accumulates noisy sensory signals over time to form a perceptual decision (Ratcliff & Smith, 2004, Roitman & Shadlen, 2002). Although pioneering work on evidence accumulation focused on the role of parietal and frontal cortices in discrimination tasks (Roitman & Shadlen, 2002; Kim & Shadlen, 1999; Gold & Shadlen, 2007), it is now clear that evidence accumulation plays a larger role in cognition and is widely distributed in the brain, spanning occipital, inferior temporal, parietal and inferior frontal cortices (Gherman et al., 2023; Goueytes et al., 2024; Imani et al., 2023; Morito & Murata, 2022; Pedersen et al., 2015; Ploran et al., 2007; Shadlen & Kiani, 2013; Tremel & Wheeler, 2015). Single neurons and neuronal populations have also been shown to accumulate evidence for detection responses mostly in the parietal cortices (Cook & Maunsell, 2002; O’Connell et al., 2012; Pereira et al., 2021). These accumulators might play a role beyond decision-making, possibly in perceptual experience (Dehaene, 2009; Kang et al., 2017; Moutard et al., 2015; Msheik et al., 2022; Pereira et al., 2021; 2022). Additionally, prior research indicates that confidence results from a readout of accumulated evidence (Desender et al., 2021; Kiani & Shadlen, 2009; Pereira et al., 2021; Pleskac & Busemeyer, 2010, van den Berg et al., 2016).

In a preregistered study (https://osf.io/8qe3a/), we used stereotactic electroencephalography (sEEG) to find neural accumulators leading to the immediate detection of weak visual stimuli. We tested whether some of these accumulators reflected conscious perception when reporting was delayed or even absent, as previous studies highlighted the importance of disentangling activity involved in the process of reporting from that involved in conscious experience per se (e.g., Cohen et al., 2020; Frässle et al., 2014; Pitts et al., 2014). Beyond stimulus detection, we identified which accumulators encoded the confidence in having detected the stimulus. Specifically, we hypothesized that confidence judgments resulted from the difference between a detection threshold and the maximal level of accumulated evidence (Pereira et al., 2022). In a nutshell, we found evidence for the existence of a neural code in the ventral visual cortex that (1) reflected evidence accumulation; (2) persisted when the report was delayed or absent; and (3) predicted participants’ confidence in seeing something.

## Results

We investigated the neural activity underlying perceptual experience, consisting of conscious access and perceptual monitoring. To this avail, we analyzed sEEG activity from 3301 recording sites in 29 human participants with drug-resistant epilepsy. In all experiments, face images were embedded for three successive frames (600 ms) at a pseudo-random point within a stream of 13 phase-scrambled images (**Figure 1A**). Face intensity was modulated to approach the participants’ detection threshold. To test whether evidence accumulation processes underlie conscious access and perceptual monitoring, we asked participants to perform three separate experiments that required different behavioral responses. We analyzed univariate and multivariate brain responses as a function of face detection, reaction times, face intensity and confidence judgments. Specifically, we looked at high gamma activity (HGA), a proxy for local neuronal firing (Lachaux et al., 2012; Leszczynski et al., 2020; Mukamel et al., 2005; Nir et al., 2007; Ray et al., 2008; Ray & Maunsell, 2011), in four pre-registered brain regions of interest (ROI): the ventral visual cortex (VVC), the superior parietal cortex (SPC), the inferior frontal cortex (IFC), and the dorsolateral prefrontal cortex (DLPFC; **Figure 1D**). Note that some intracranial electrodes targeting the IFC had their deepest channels in the anterior insular cortex. For simplicity, we will refer to the IFC as a ROI, and will focus on the anterior insula when appropriate.

**Figure 1.**
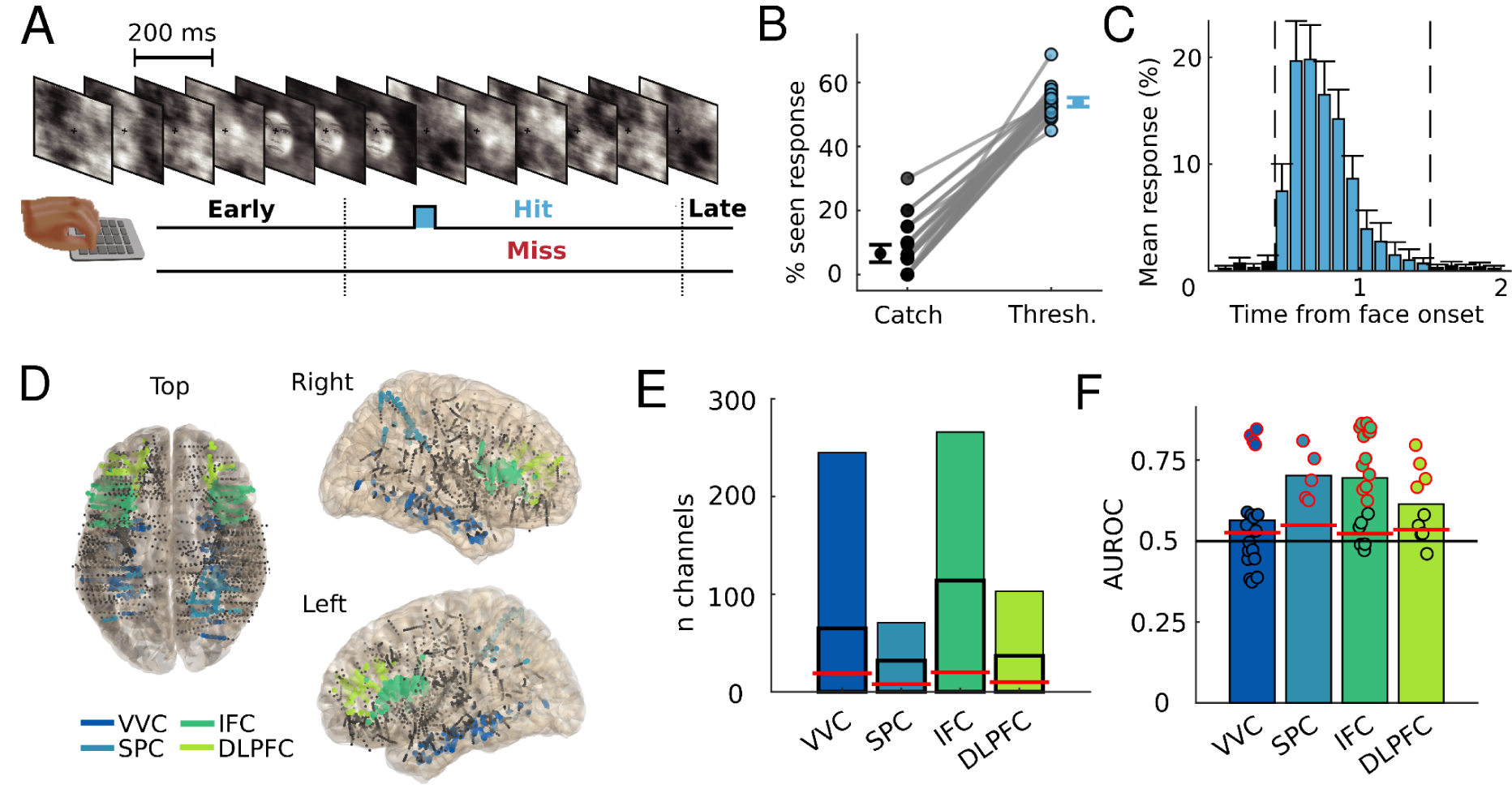
Experiment 1. *A.* Experimental procedure. Participants fixated on a cross superimposed on scrambled frame images. An embedded face stimulus was presented for three consecutive 200 ms frames at a random time starting 600 ms to 1800 ms after trial onset. Face stimuli do not represent actual people: they are AI-generated and freely available for scientific purposes on this website: https://generated.photos/. When participants pressed the button between 400 ms and 1500 ms after face stimulus onset, the trial was considered a hit (trials with response times < 400 ms or > 1500 ms were excluded, see panel C and methods). When participants did not press, the trial was considered a miss. *B.* Average and individual detection rates for all participants, both in catch trials where no target was presented and in threshold trials where the target was presented at participants’ detection threshold. *C.* Mean percentage of responses as a function of reaction times (100 ms bins) across participants. Dashed vertical lines show the 400 ms and 1500 ms limits for considering the trial as a hit. *D.* Analyzed channels normalized to an MNI template, with channels in the ventral visual cortex (VVC), superior parietal cortex (SPC), inferior frontal cortex (IFC) and dorsolateral prefrontal cortex (DLPFC) in color. *E.* Total number of channels in the four ROIs. The box with black contours shows the proportion of responsive channels, defined as channels showing a significant positive difference between hit and miss trials. The red line indicates the maximum number of responsive channels expected by chance (α = 0.05). *F.* Cross-validated average area under the receiving operator characteristic (AUROC) for the hit vs. miss decoder in the four regions of interest (bar plot: average; dots: individual participants). Red lines indicate group-level chance-level performance (assessed by permuting labels across trials). Red dot surrounds participants who displayed above chance-level performance after false discovery rate correction. All error bars indicate 95% confidence interval.

### Evidence accumulation activity is found in the ventral visual and inferior frontal cortices

We started by establishing which brain channels and regions reflected immediate detection and evidence accumulation for such detection. In experiment 1, participants provided an immediate report - they were instructed to press a key as soon as they detected a face. Trials were considered a hit if participants pressed in a window finishing 1500 ms after face onset (53.86±3.99%; mean±SD; **Figure 1B-C**), and a miss if they did not press. We removed trials with premature responses (up to 400 ms after the face onset; 2.16±2.32%). We identified responsive channels (n = 1002, 30.08±11.88% of channels across participants, p < 10^−5^, binomial test), defined as showing more HGA in hit than miss trials during the second following face onset using nonparametric clustering analyses (Maris & Oostenveld, 2007). Responsive channels were found in significant numbers in the four ROIs (all ps < 10^−5^, **Figure 1E**, see **Supplementary Figure 1** for other brain areas). The number of channels with more HGA in miss than hit trials was not above the level expected by chance, either across all channels (n = 158, 4.79%, p = 0.7) or in ROIs (all ps > 0.28). Hierarchical mixed-effects regressions fitted by time point and corrected for false discovery rate indicated that the time courses for the effect of immediate detection differed qualitatively across ROIs (0.31 to 0.94 s after face onset in the VVC, 0.06 to 0.9 s in the SPC, 0.28 to 1 s in the IFC and 0.44 to 1 s in the DLPFC). To characterize the latent variable leading to immediate detection, we developed a time-resolved decoding algorithm that classifies a trial as a hit when the maximum value of the weighted sum of all channels in the region of interest reaches a threshold, similar to a decision variable within an evidence accumulation framework (Shadlen & Kiani, 2013). The decoders could successively classify hit vs. miss trials in all four ROIs (**Figure 1F**). Further multivariate analyses are focused on those participants for which decoding was successful (VVC: 4/18 participants; SPC: 5/5; IFC: 13/19; DLPFC: 4/9; **Supplementary table 3**). To summarize, univariate and multivariate analyses indicated the implication of several brain areas during immediate stimulus detection in experiment 1, including our four pre-registered ROIs.

To specify the role of evidence accumulation in stimulus detection, we examined the relationship between neural activity and detection times in hit trials. More precisely, we sought to identify two neural markers of evidence accumulation. First, we looked for a hallmark of evidence accumulation: A negative correlation between response times and the derivative of stimulus-locked HGA, underlying the fact that faster responses are associated with a steeper slope of HGA following stimulus onset (Cook & Maunsell, 2002; O’Connell et al., 2012). Such an effect of reaction times on the slopes of HGA was observable in 18.46% of all responsive channels (p < 10^−5^, binomial test, **Figure 2A** shows examples of such channels). At the level of ROIs, we found an effect of reaction times on the slope of HGA in the VVC (t(2427) = 4.72; p < 10^−5^) and the IFC (t(5465) = 3.83; p = 1.3 x 10^−4^; **Figure 2B**), but not in the SPC (t(1464) = 1.86; p = 0.06) nor in the DLPFC (t(1810) = 1.38; p = 0.17; **Supplementary Figures 2A and 3**). Second, we investigated whether the aforementioned latent variable trained to discriminate hit vs. miss trials (see above) also reflected evidence accumulation. We tested if the timing at which the decoded latent variable crossed a detection threshold (set individually for each participant to balance hit and miss trial detection) would predict response times. (**Figure 2C**). A positive correlation between reaction times and the timing of threshold crossing by the latent variable was found in all ROIs: for all 4 participants in the VVC, for 7 out of 13 in the IFC and for 2 participants out of 5 in the SPC and for one out of 4 the DLPFC (**Figure 2D-E; Supplementary table 3**). Overall, these findings provide converging evidence that the VCC and IFC instantiate evidence accumulation pertaining to stimulus detection.

**Figure 2.**
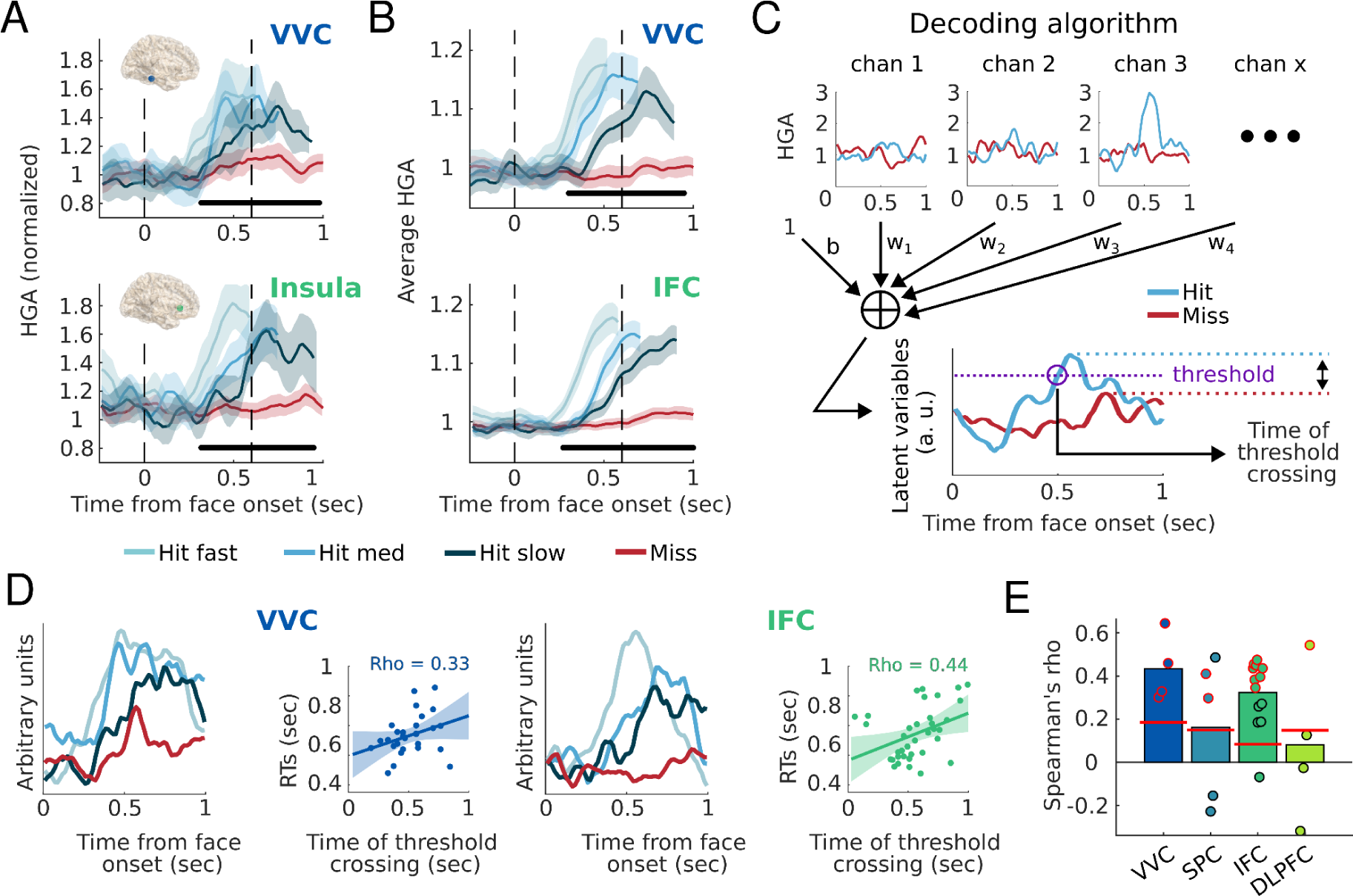
Evidence accumulation signals. *A.* HGA traces from single channels in the VVC (top) and IFC (bottom). HGA in miss trials (red traces) and in hit trials separated in three terciles based on reaction times (blue traces). The vertical dashed lines indicate the onset and offset times of the face stimuli. The black bars indicate significant effects of detection (cluster-based permutation tests). Note that the channel in the insula was localized by the atlas as part of the IFC. *B.* Averaged HGA across all responsive channels in the VVC (top) and IFC (bottom).The black bars indicate main effects of detection as assessed with sample-by-sample hierarchical mixed-effects regressions (corrected for false discovery rate across time). Response-locked plots can be found in **Supplementary Figure 4**. *C.* Representation of the decoding algorithm, with example hit and miss trials. A trial-by-trial decision signal was obtained by applying the weights of the multivariate classifier on each time point. The weights were chosen to optimize the distance between the maximum values of the latent variables in hit vs. miss trials. *D.* Latent variables decoded from the VVC of one participant (G5; left panel) and the IFC of another participant (G21; right panel), averaged over all miss trials and separated in three terciles based on reaction times for hit trials. Insets: Trial-by-trial depiction of the relationship between reaction times and the predicted time of threshold crossing of the decoded latent variable, with Spearman correlations. *E.* Average Spearman correlation between observed and decoded response times across participants (bar plot: average; dots: individual participants). Red lines indicate group-level chance-level performance assessed by permuting labels across trials. Red dot surrounds indicate participants who displayed above chance-level performance after false discovery rate correction. All shaded areas represent 95% confidence intervals.

In experiment 1, we identified two regions that accumulated evidence when participants were required to respond immediately after detecting the stimulus. It remained unclear whether evidence accumulation in these regions was associated with conscious access per se, or rather with decision-making and motor processes. We addressed this question in two further experiments.

### The perceptual decision signal decodes delayed detection and passive perception

Immediately after completing experiment 1, participants performed a delayed-report (experiment 2) and a no-report (experiment 3) version of the task on the same stimulus sequence (**Figure 3A**). In experiment 2, signals related to perceptual decisions but free of motor confounds were obtained by asking participants to wait for the end of the trial (i.e., a minimum of 1300 ms after stimulus onset) to provide a detection response. Face stimuli were either presented at perceptual threshold (*threshold trials*; 47.34±7.04% trials reported as seen) or at a higher intensity defined as 125% the perceptual threshold (*suprathreshold trials*, 76.5±10.27% trials reported as seen, significantly more than in threshold trials; t = 18.07, p < 10^−5^; **Figure 3B and Supplementary Figure 5**). In 17% of trials, no face was presented at all (*catch trials*). Participants could report that they saw more than one face in cases where a non-veridical percept of a face occurred during the same trials as a veridical percept of the stimulus. To make this response option seem sensible to the participants, we included one trial in each block that actually contained two suprathreshold faces, but excluded them from analyses. In experiment 3, we tested whether perceptual signals from the previous experiments were still present in the absence of any task, during passive viewing of face stimuli at threshold and suprathreshold intensities.

**Figure 3.**
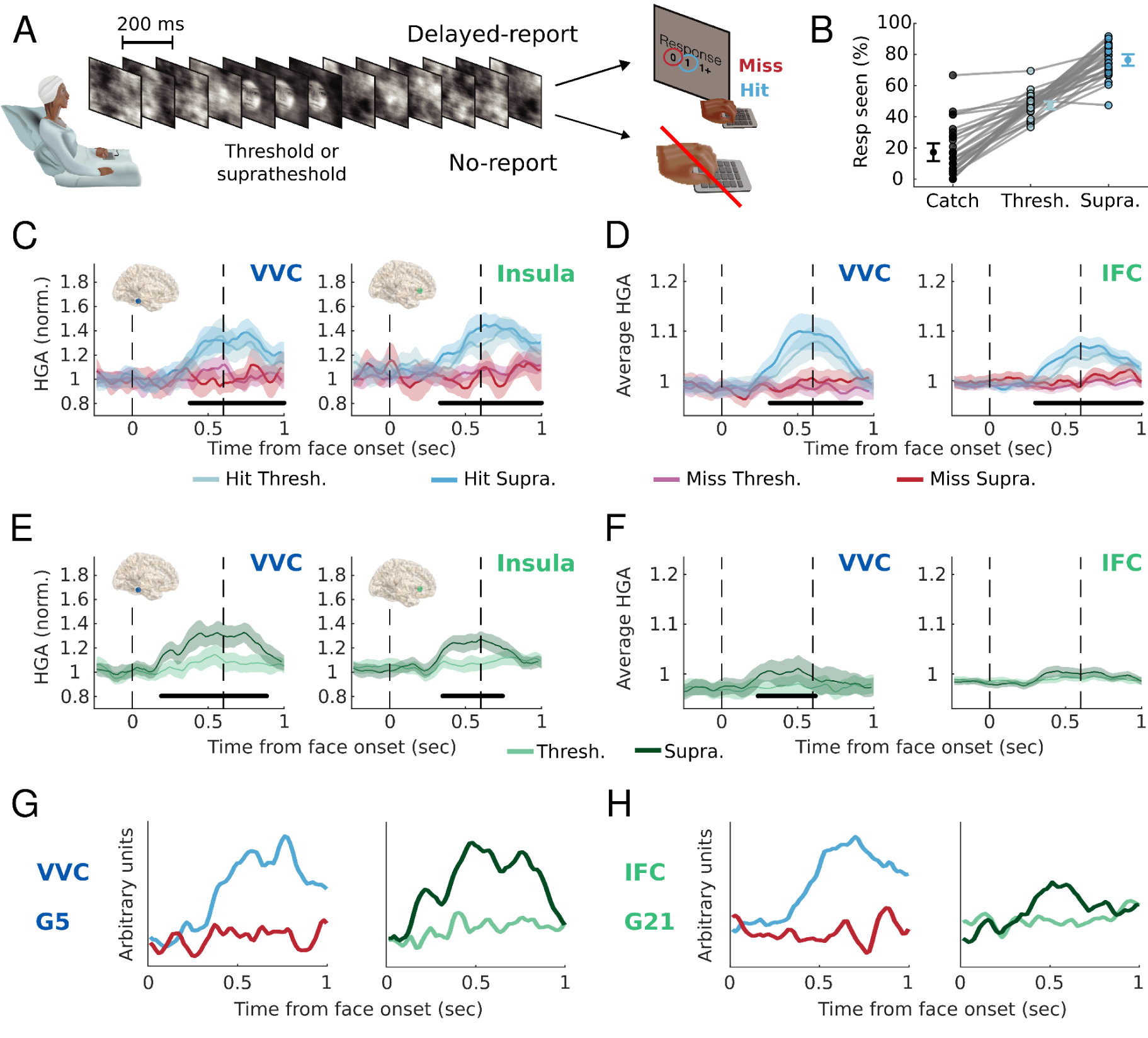
Experiments 2 and 3. *A.* Experimental procedures. In experiment 2, participants reported a detection response at the end of the trial. In experiment 3, participants passively viewed the visual stream. *B.* Mean and individual false alarm rates in catch trials and hit rates when faces were presented at threshold and suprathreshold intensities in experiment 2. *C.* Averaged HGA traces from the same channels as in **Figure 2A**, as a function of stimulus detection and intensity in experiment 2. *D.* The averaged activity across all responsive channels in the VVC (left panel) and IFC (right panel) in experiment 2. *E.* Averaged HGA traces from the same channels as in **Figures 2A and 3C**, as a function of stimulus intensity in experiment 3 (right panels). *F.* Similarly to *D*, the averaged activity across all responsive channels in the VVC (left panel) and IFC (right panel) in experiment 3. In *C.* and *D.*, the black bars indicate significant main effects of detection; in *E.* and *F*., they indicate significant main effects of stimulus intensity (cluster-based statistics for individual channels, hierarchical mixed-effects regressions for ROIs; corrected for false discovery rate across time). *G.* Averaged decoded latent variable in hit (blue trace) and miss (red trace) trials in experiment 2 (left panel) and in threshold and suprathreshold trials in experiment 3 (right panel) in the VVC of participant G5. *H.* Same as *G* in the IFC of participant G21. All error bars and shaded areas indicate 95% confidence intervals.

Over a third of channels identified as responsive in experiment 1 also reflected detection in the absence of motor preparation in experiment 2 (39.22%, p < 10^−5^, binomial test). This number increased to over half when looking only at channels that reflected evidence accumulation in experiment 1 (57.3%; see examples in **Figure 3C**). Hierarchical mixed-effects regressions fitted to the average value of HGA in the responsive time window (see *Methods*) with detection reports and stimulus intensity as predictors showed a main effect of detection in all ROIs (VVC: t(7953) = 5, p < 10^−5^; SPC: t(3832) = 3.5, p = 4.73 x 10^−4^; IFC: t(13722) = 4.53, p < 10^−5^; DLPFC: t(4520) = 2.62, p = 0.0089, **Supplementary Figures 2B and 6A**; see also **Supplementary Figure 7** for a control analysis). HGA was also influenced by stimulus intensity in the VVC (t(7953) = 3.47, p = 6.15 x 10^−4^), the SPC (t(3832) = 2.88, p = 0.004), the IFC (t(13722) = 5.09, p < 10^−5^), but not in the DLPFC (t(4520) = 1.94, p = 0.053; **Figure 3D and Supplementary Figure 6C**). There was a significant interaction between detection and stimulus intensity in the VVC (t(7953) = 2.76, p = 0.006) and IFC (t(13722) = 2.39, p = 0.017; **Supplementary Figure 6E**), which was explained by stimulus intensity influencing HGA in hit trials only. Time-resolved hierarchical mixed-effects regressions indicated that the effects of detection in experiment 2 occurred for similar durations as immediate detection effects in experiment 1 (0.33 to 0.91 s after face onset in the VVC, 0.46 to 1 s in the SPC, 0.31 to 1 s in the IFG and 0.67 to 0.96 s in the DLPFC), and that the interaction effect started around that same time in the VVC (0.34 s) while it emerged later in the IFC (0.57 sec). Thus, all ROIs encoded stimulus detection irrespective of motor responses and stimulus intensity.

These results from experiment 2 do not help arbitrate whether regions only played a role in taking and maintaining a decision or in conscious access itself. We reasoned that for ROIs solely involved in decision-making, passive viewing in experiment 3 would not cause stimulus-locked HGA activity. On the contrary, ROIs involved in conscious access irrespective of report should have higher HGA in trials where faces were presented at suprathreshold (defined as 150% the perceptual threshold) than threshold intensity. This is because faces would have been perceived more often in suprathreshold trials even during passing viewing (i.e., a no-report condition). In experiment 3, we found this effect of stimulus intensity in 9.68% of all responsive channels (p < 10^−5^, binomial test), and in 22.64% of responsive channels that reflected evidence accumulation in experiment 1 and detection in experiment 2 (examples in **Figure 3E**), but only in 4% of non-responsive channels (p = 0.99). Hierarchical mixed-effects regressions showed that HGA was influenced by stimulus intensity in the VVC (t(5750) = 3.27, p = 0.001), but did not show this effect in other ROIs (SPC: t(2296) = 0.66, p = 0.51; IFC: t(13720) = 1.83, p = 0.067; DLPFC: t(4623) = −0.24, p = 0.98; **Figure 3F and Supplementary Figures 2C and 8A**). Several channels did show responses in all experiments in the anterior insula (**Figure 4B**), although an effect of stimulus intensity in experiment 3 was not observed at the level of the region. Interestingly, the only other region to show an effect of stimulus intensity in exploratory whole brain analyses was the lateral visual cortex adjacent to the VVC, despite many regions showing an effect of detection in experiment 2 (**Supplementary Figures 6B and 8B**). The effect of stimulus intensity in the VVC started earlier (0.25 to 0.61 s after face onset) than effects of detection in experiment 1 and 2. Overall, these results indicate that while many regions were sensitive to detection without motor responses, only the ventral and lateral visual cortices were also responsive to stimulus perception in the absence of overt reports.

**Figure 4.**
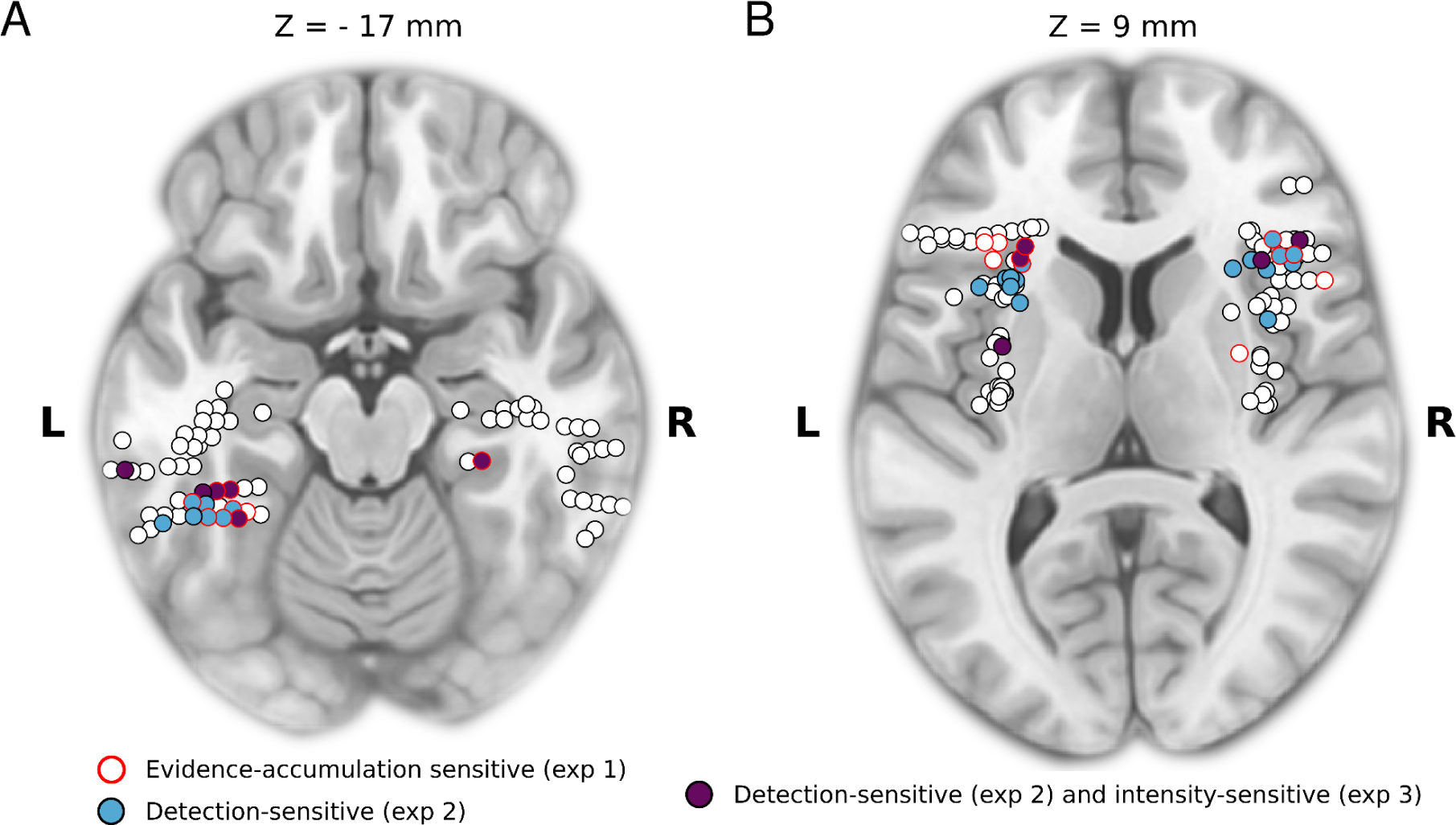
Channel locations. *A.* Projection of channels localized in the VVC and lateral visual cortex on a transverse plane of the MNI template brain at coordinate Z = −17 mm (see **Supplementary figures 9-10** for other slides and localizations in native space). *B.* Same projection for channels in the IFC and insula at Z = 9 mm. Channels within 2 mm of the displayed slice are represented. The responsive channels reflecting evidence accumulation in experiment 1 are circled in red, those showing a main effect of detection in experiment 2 are colored in blue and those showing this effect and an effect of stimulus intensity in experiment 3 are colored in purple. Note that many channels that were localized in the IFC by our automatic pipeline are actually located in the anterior insular cortex. A control analysis following a visual inspection by a trained neurologist revealed similar results in the anterior insula and in our pre-registered IFC ROI (**Supplementary figure 11**).

We showed that activity in the VVC is consistent with evidence accumulation and persists when reporting is delayed or even absent. However, these results could be driven by different neural populations within the same brain region that accumulate evidence towards a motor response in experiment 1, delayed-report in experiment 2, or passive viewing in experiment 3. We accordingly tested whether a common neural population instantiating evidence accumulation as shown in **Figure 2** was involved in all three experiments. Using the weights from the multivariate decoders trained to discriminate between hits and misses in experiment 1, we decoded delayed stimulus detection in experiment 2 in all regions (VVC: 4/4 participants showing this effect; SPC: 3/5 participants; IFC: 4/13 participants; DLPFC: 3/4 participants; **Figure 3G-H; Supplementary table 3**). We then used a similar approach to decode face intensity in experiment 3. Suprathreshold vs. threshold trials were used as labels under the premise that activity in suprathreshold trials would resemble activity in hit trials in the previous two experiments. Face perception could be decoded with high performance in the VVC (AUROC = 0.69) for one out of two participants who performed experiment 3 (for the remaining two participants we could not acquire recordings in this experiment due to clinical constraints; **Figure 3G**). No significant decoding was found for any participant in the other ROIs (**Figure 3H; Supplementary table 3**). Interestingly, high decoding performance was observed in the VVC despite the fact that some suprathreshold stimuli may have remained unperceived in some trials, and that threshold stimuli were arguably perceived in about half the trials. In line with our univariate results, stimulus intensity could also be decoded for one participant in the lateral visual cortex (**Supplementary table 4**). When taking all the above results into account, the same latent variable in the VVC reflected immediate detection and evidence accumulation in experiment 1, delayed detection in experiment 2, and stimulus intensity during passive viewing in experiment 3. In other words, we propose that noisy sensory information is accumulated in the VVC even in the absence of task demands. Interestingly, the shared response across experiments identified in the VVC appeared to be driven by channels located at a similar location in the middle to anterior fusiform gyrus (**Figure 4A and Supplementary figure 9**).

Until now, we focused on conscious contents that were driven by external sensory stimuli. In pre-registered analyses, we next assessed whether the latent variable which reflected conscious sensory percepts in the VVC could also reflect non-veridical percepts, in trials where participants reported seeing a face when none was presented on the screen. As we had no external event to time-lock our analysis to, we examined latent variables in the entire trial window. Doing so, we found that decoded latent variables crossed a decision threshold more often in false alarm trials (17.19±15.86% of trials with no face) than in correct rejection trials (p = 0.034, permutation test; **Figure 5a**) in experiment 2. Moreover, we found that this threshold was also crossed more often when participants reported seeing more than one face (6.87±7.91% of trials where one face was presented) vs. only one face (p = 0.023, permutation test; **Figure 5b**). Neither of those effects was found in any of the other three ROIs or in the lateral visual cortex. These results, combined with the results of passive viewing in experiment 3, indicate that the neural code identified in the VVC reflects both veridical and non-veridical conscious percepts.

**Figure 5.**
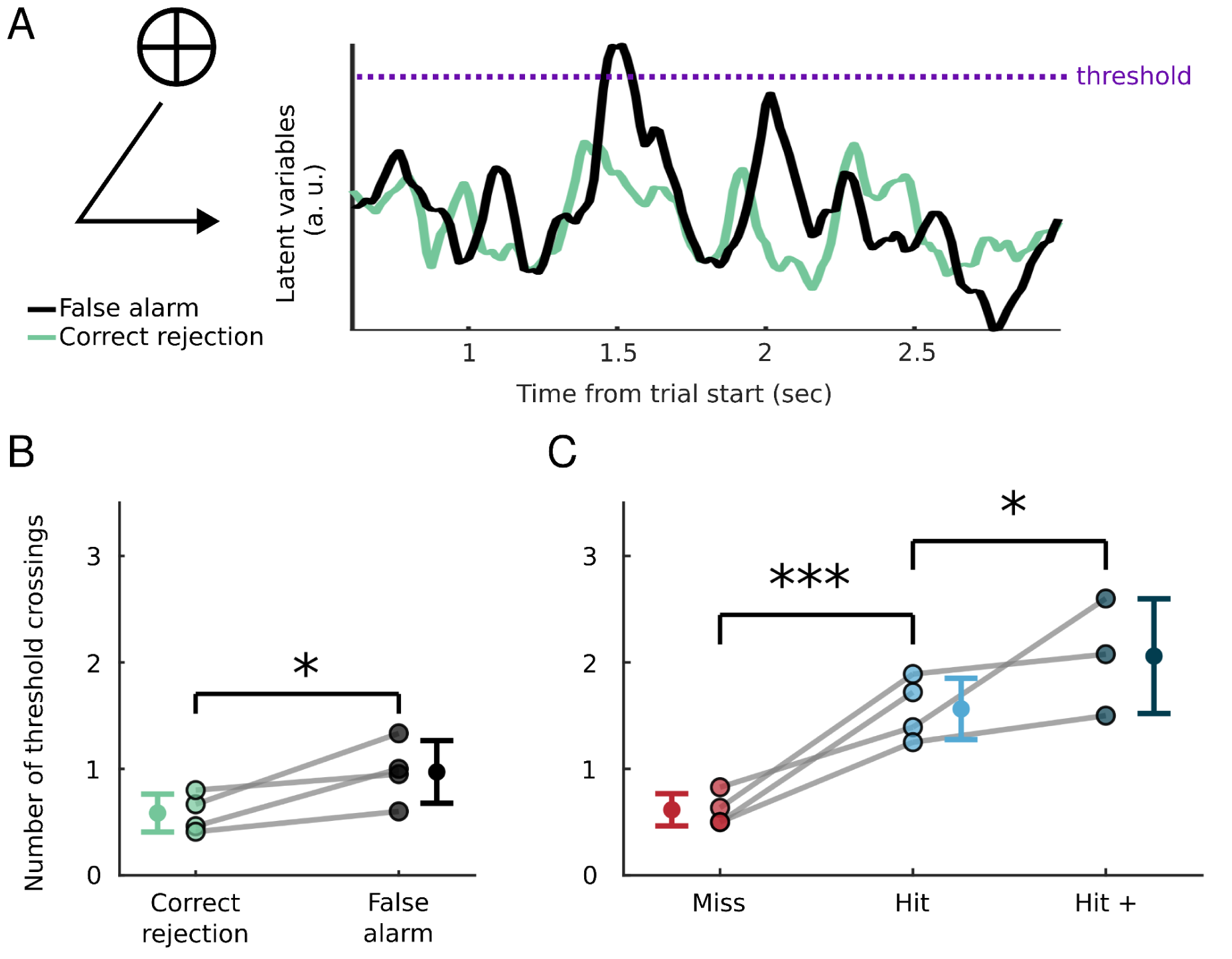
Non-veridical percepts in the VVC in experiment 2. *A.* The weights from the decoding algorithm represented on **Figure 2C** were applied to the entire trial window in which no face was presented. The traces are examples of the decoded latent variable for false alarm (black) and correct rejection (green) trials. We measured the number of times that the variables crossed a decision threshold, set individually for each participant to balance hit and miss trial detection. *B.* Participant and overall averages of the number of times that the decoded latent variable crossed the detection threshold, for trials where no face was presented and the participant reported seeing no face (correct rejection) vs. one face (false alarm). *C.* The same for trials where one face stimulus was presented and the participant reported seeing no face (miss) vs. one face (hit) vs. more than one face (hit +). The results are for all participants that showed significant decoding of hit vs. miss trials in the stimulus onset-locked window in experiment 1. Missing values correspond to participants without trials in one response category. P-values were obtained with permutations performed on the relevant trials for each participant and then comparing the participants’ average difference between the two conditions of interest to the permuted distribution of this difference: *** = p < 0.001, ** = p < 0.01, * = p < 0.05. Error bars indicate 95% confidence intervals.

In this section, we have shown that the same neural code is involved in evidence accumulation and conscious access, one aspect of perceptual experience. In the next section, we ask whether another aspect of experience, perceptual monitoring, also relies on this same neural code.

### Evidence accumulation determines confidence in the ventral visual cortex

In experiment 2, participants were probed for their confidence in their detection response (**Figure 6A**). A hierarchical mixed-effects regression on confidence revealed no significant main effect of stimulus intensity (t = 0.83, p = 0.41), but a main effect of detection (t = 2.53, p = 0.01) and a significant interaction between stimulus intensity and detection (t = 4.88, p < 10^−5^). To explore this interaction further, we analyzed hit and miss trials separately, and found that confidence increased with stimulus intensity for hit (t = 7.14, p < 10^−5^), but not for miss trials (t = 0.13, p = 0.9; **Figure 6B**). These results indicate that face intensity influenced confidence reports in hit trials only, consistent with limited confidence sensitivity when reporting on the absence of a stimulus (Mazor et al., 2020; Meuwese et al., 2014; Pereira et al., 2021).

**Figure 6.**
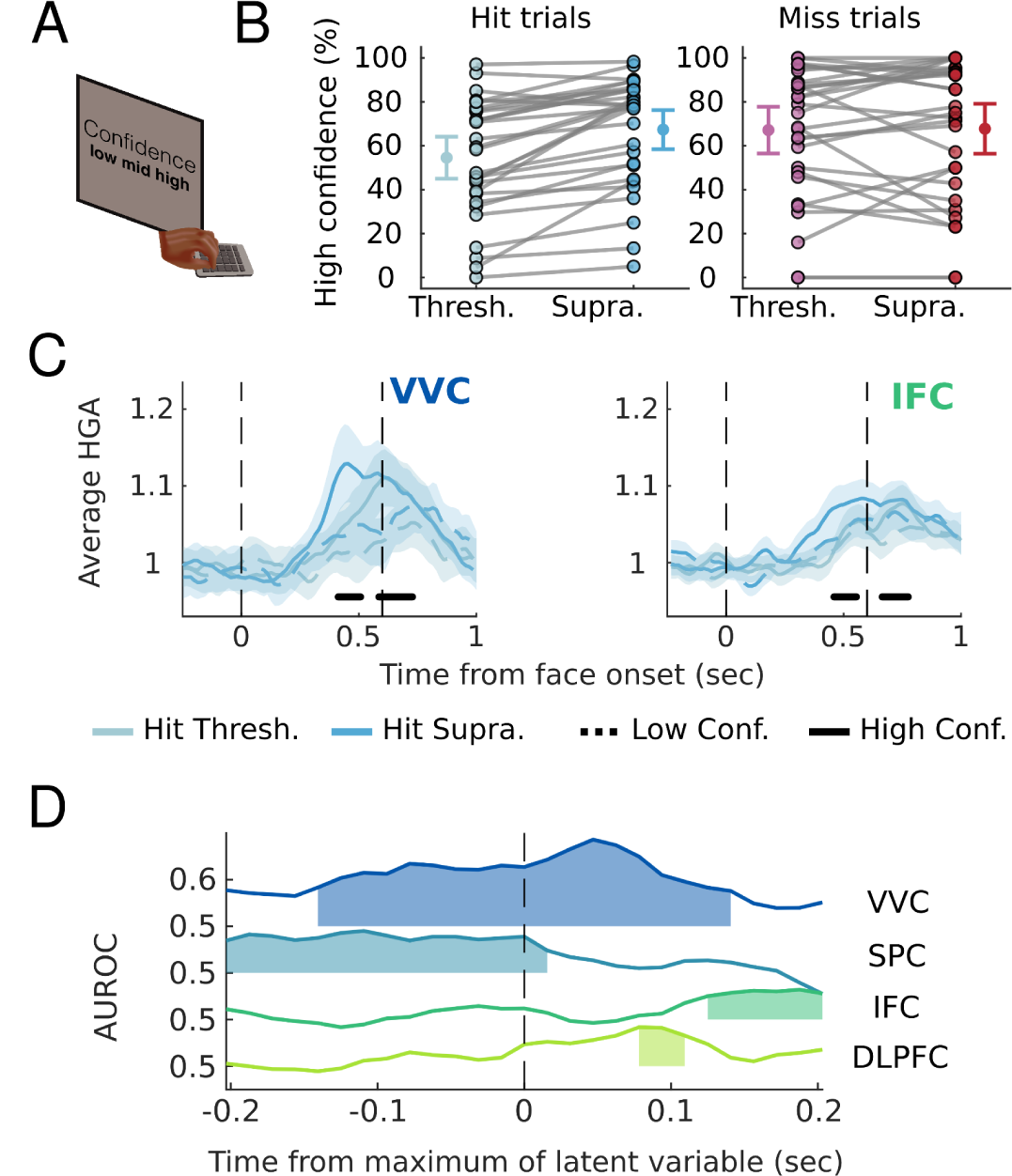
Confidence judgments in experiment 2. *A.* participants were probed on a 3-point scale for their decision confidence after providing the detection response. Confidence reports were binarized by pooling “moderately sure” and “unsure” responses as “low confidence” and reporting “very sure” responses as “high confidence” (see **Supplementary Figure 5B**). *B.* Mean and individual percentage of high confidence trials as a function of stimulus intensity (threshold vs. suprathreshold) for hit trials (left panel) and miss trials (right panel). *C.* Averaged HGA traces from hit trials in one channel in the VVC (left panel) and one channel in the IFC (right panel) as a function of confidence and stimulus intensity. The black bars indicate significant clusters for the effect of confidence. *D.* The averaged activity for hit trials as a function of confidence and stimulus intensity across all responsive channels in the VVC (left panel) and IFC (right panel). The black bars indicate significant main effects of confidence (corrected for false discovery rate across time). *E.* AUROC for decoding high vs. low confidence in hit trials, as a function of time around the maximum of the decoded latent variable. All error bars and shaded areas indicate 95% confidence intervals, except in E where shaded areas indicate above chance-level performance at the group level (assessed by permuting labels across trials, uncorrected).

We investigated whether brain activity in experiment 2 reflected confidence. Some responsive channels (13.07%, p < 10^−5^, binomial test) displayed more HGA in high confidence than low confidence hit trials. Hierarchical mixed-effects regressions with confidence and stimulus intensity as predictors showed that higher confidence was associated with more HGA in three ROIs (VVC: t(4862) = 2.44, p = 0.015; SPC: t(2346) = 2.86, p = 0.004; IFC: t(8615) = 3.88, p = 1.07 x 10^−4^; **Figure 6C**) but not in the DLPFC (t(2804) = 0.98, p = 0.33, see **Supplementary figures 3D and 12A**). In miss trials, no effect of confidence was found in any of the regions (VVC: t(3081) = 0.72, p = 0.47; SPC: t(1475) = 0.86, p = 0.39; IFC: t(5107) = 1.64, p = 0.1; DLPFC: t(1716) = −0.15, p = 0.88, see **Supplementary figures 12C**). These results indicate that several ROIs, including the VVC, encoded confidence in seeing a face. The effect of confidence appeared to be reflected in a higher maximum of HGA, while stimulus intensity modulated the slope of the HGA increase (**Figure 6C**), consistent with a recent proposal that confidence is computed as the difference between the maximal accumulated evidence and the detection threshold (Pereira et al., 2021; 2022).

We next investigated whether the multivariate decision signal identified in experiment 1 could also explain confidence judgments. To characterize the temporal dynamics with which confidence is built, we assessed how decoders predicted participants’ confidence in having seen a face before, around and after the maximum of the decoded latent variable (**Figure 6D**). In the VVC, decoding performance across participants was significant during a window of ∼300 ms centered on the maximum, and then went back down to chance-level. These findings support the hypothesis that confidence judgments result from the difference between a detection threshold and the maximal level of accumulated evidence (Pereira et al., 2021; 2022). By contrast, confidence was better predicted before maximal evidence was reached in the SPC, and after this point in the IFC and DLPFC.

### A leaky evidence accumulation process reproduces the neural activity

Our empirical results converge to indicate that the VVC accumulates evidence pertaining to conscious access and perceptual monitoring, irrespective of reports. To more formally establish the role of evidence accumulation for conscious access, we assessed if a computational model based on a leaky evidence accumulation process could reproduce both the behavioral and neural data we observed (Pereira et al., 2021; 2022). Compared to standard accumulation models, the model included a leakage term which circumvents the continuous accumulation of noise in a detection task with uncertain timings of stimulus onset (Cook & Maunsell, 2002; Ossmy et al., 2012; Usher & McClelland, 2001; see *Methods* for more details). We simulated 10000 traces of evidence accumulation and the corresponding detection responses and response times. We could thereby reproduce not only the behavioral results (**Supplementary figure 13**) but also the neural data (**Figure 7B**) which resembled several idiosyncratic features of the HGA responses observed in the VVC (**Figure 7A**). These include the slightly lower maximum observed for slower hit trials in experiment 1, as well as the effect of intensity in hit trials but not in miss trials in experiment 2. Model simulations therefore support the role of leaky evidence accumulation in stimulus detection, and suggest that this computational mechanism may be implemented in the VVC.

**Figure 7.**
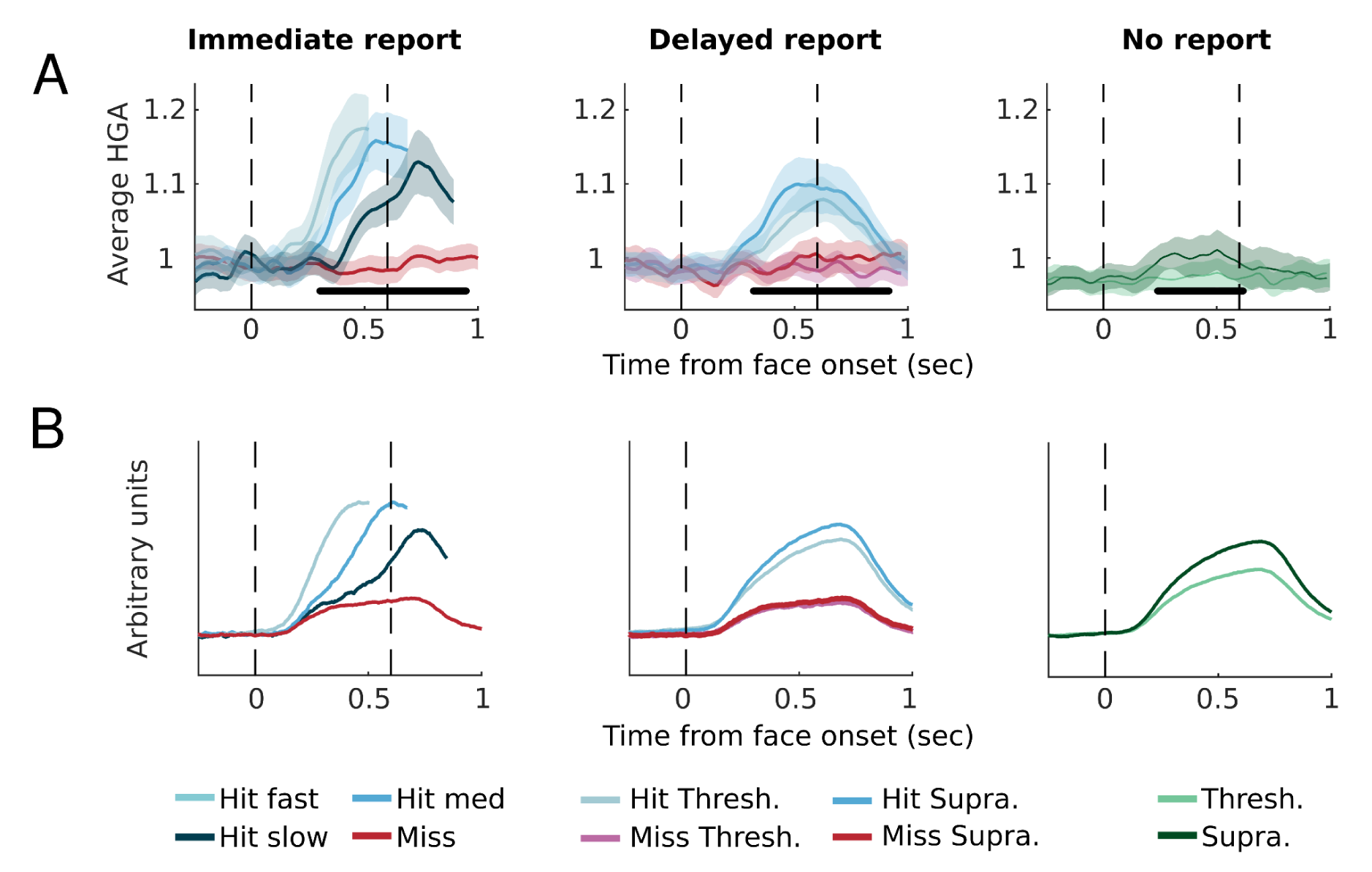
Model simulations reflect HGA in the VVC. *A.* Averaged activity in the VVC in the three experiments, reproduced from **Figure 2B and Figure 3D-F** for comparison with the simulation results. *B.* The corresponding simulation traces averaged across repetitions. Traces are plotted as a function of detection and reaction times in experiment 1 (left panel), of detection and intensity in experiment 2 (middle panel) and of intensity in experiment 3 (right panel).

## Discussion

We confirmed that neural activity in the VVC verified three pre-registered hypotheses linking it to perceptual evidence: it relates to evidence accumulation in experiment 1, discriminates hits from misses in experiment 2 and is modulated by the stimulus in experiment 3. Amongst our four pre-registered ROIs, both the VVC and IFC were found to accumulate evidence for face detection in experiment 1, as shown by two markers of evidence accumulation. First, reaction times explained the slope of the HGA in responsive channels in these regions. Second, reaction times also correlated with the time at which an estimated latent variable, tuned to distinguish seen and unseen faces, crossed a threshold. Hierarchical mixed-effects regressions applied to responsive channels showed that while activity in all ROIs discriminated between hit and miss trials in experiment 2, only the VVC was influenced by stimulus intensity during passive viewing in experiment 3. This result was further substantiated by multivariate analyses showing that the signal underlying detection-sensitivity in the VVC in experiment 1 could generalize to delayed (experiment 2), passive (experiment 3) and non-veridical face perception (experiment 2). Finally, univariate analyses indicated that participants’ confidence in having seen a face was encoded in the VVC, SPC and IFC. Interestingly, our decoding algorithms trained to distinguish between hit and miss trials in experiment 1 could decode confidence judgments at the time of maximal accumulated evidence in hit trials in experiment 2 in the VVC and SPC, but not in the IFC. We note that these results are likely to be valid in the general population, given that sEEG recordings concern mostly healthy cortical tissue (Parvizi & Kastner 2018).

### Widespread evidence accumulation indicates different functional roles of accumulators in the brain

First, our results bring new insights regarding the functional roles of neural accumulators and about their interactions (DaPasquale et al., 2021; Hanks et al., 2015; Morito & Murata, 2022; Msheik et al., 2022; O’Connell & Kelly, 2021). Previous fMRI (Morito & Murata, 2022; Pedersen et al., 2015; Ploran et al., 2007; Tremel & Wheeler, 2015) and intracranial EEG (Gherman et al., 2023; Goueytes et al., 2024; Liu et al., 2023) research uncovered widely distributed accumulators in discrimination tasks with different activity profiles (**Supplementary Figure 4**). Our results show that cortical evidence accumulation is also widespread and diverse for a detection task. Exploratory analyses on all brain areas pointed to six regions accumulating evidence apart from the IFC and the VVC: the lateral visual cortex, the insula, the dorsolateral motor cortex, the dorsolateral premotor cortex, the ventral somatosensory cortex and the ventrolateral orbitofrontal cortex (**Supplementary figure 3**). All regions reflecting evidence accumulation were also found to reflect stimulus detection when the report was delayed in experiment 2, indicating that they were not merely involved in the preparation of a motor action. In previous studies, experimental manipulations (e.g., temporal delay, change of effector) made it possible to isolate effector-independent accumulators in the brain, displaying accumulation for abstract decisions (e.g., Gherman et al., 2023; Twomey et al., 2016; Pereira et al., 2020). Going one step further, we tested whether accumulators were involved in perceptual processes irrespective of a decision. Previous works using scalp EEG have shown that centro-parietal markers of evidence accumulation thought to reflect subjective experience (Tagliabue et al., 2019) vanish during passive viewing (Twomey et al., 2016; Cohen et al., 2020; Pitts et al., 2014). In line with these results, for most regions in our study, neural signatures of stimulus detection also vanished in experiment 3. Crucially, however, in the ventral and lateral visual cortices, we found stimulus-locked activity during passive viewing with similar timings as evidence accumulation activity during stimulus detection. Our results thus suggest that evidence accumulation in these regions persists irrespective of the preparation of a report, indicating that it might be involved in conscious access (Aru et al., 2012; Frässle et al., 2014; Sergent & Naccache, 2012; Tsuchiya et al., 2015). Scalp EEG studies may fail to capture the activity that we observe in the VVC due to its anatomical localization below the temporal lobe (see below).

### Evidence accumulation for conscious access was most evident in the mid-fusiform gyrus

Because of the uncontrolled and sparse sampling of sEEG, the absence of evidence for neural signatures of evidence accumulation and perceptual experience in other ROIs than the VVC should not be interpreted as conclusive evidence for their absence (Parvizi & Kastner, 2018). This is particularly problematic in the SPC, as we could not collect data in experiment 3 for two participants who showed decoding of delayed detection in experiment 2 in this region (**Supplementary table 3**). Previous studies showed that neurons in the SPC accumulated evidence in a diversity of tasks (Kiani & Shadlen, 2013; Pereira et al., 2021) and encoded the intensity of vibrotactile stimuli in a no-report task (Pereira et al., 2021). In the IFC, although 7 participants were found to show evidence accumulation, none of the decoders could generalize to the passive viewing condition in experiment 3 despite some single channels showing a strong effect of stimulus intensity (see **Figure 3E**). Note that these channels appeared to be specifically located in the anterior insular cortex at the junction with the frontal operculum (see **Figure 4B and Supplementary figure 9**), an area that was responsive to conscious access in previous no-report studies (Dellert et al., 2021; Weilnhammer et al., 2021). There is a possibility that decoding did not transfer from experiment 1 to experiment 3 in this region due to the sparse sampling of sEEG. Alternatively, our decoders may have failed to generalize from experiment 1 to experiment 3 because of the algorithm they were based on. This algorithm assumes that conscious perception is reflected by the maximum amount of accumulated evidence, while insular and frontal evidence accumulation processes might reflect report at the macroscopic level and not consciousness per se (but see Kapoor et al., 2022). Decoding transfer was observed in the VVC, and appeared to be driven by channels located around the mid-fusiform gyrus (**Figure 4A and Supplementary figure 9**). This cortical location, which include the fusiform face area, has been found to be responsive to faces using several imaging methods (Kanwisher et al., 1997; Jonas et al., 2016; Khuvis et al., 2021; Rossion et al., 2023), to be involved in the recall of faces (Khuvis et al., 2021), and to lead to distorted perception of faces when stimulated (Parvizi et al., 2012; Schrouff et al., 2021). This specificity can explain why stimulus detection in experiment 1 was decoded in only 4 participants out of 17 in the VVC, all of whom showed high decoding accuracy (AUROC ∼80%) and reflected evidence accumulation. Notably, all those participants had channels located in the mid-fusiform gyrus. Because of clinical constraints, we only had data for two out of these four participants (G5 and G18) in experiment 3. Participant G5, but not G18, showed successful decoding of stimulus intensity in experiment 3 (note again that decoding performance for this participant was high, AUROC = 0.69). We see at least two possible explanations for the differences in decoding success across these participants. On the one hand, evidence accumulation for conscious access might be performed by some parts of the responsive visual cortex and not others. On the other hand, some participants might have had lower evidence accumulation rates or higher perceptual thresholds during passive viewing in experiment 3, and therefore failed to see most faces. Anecdotally, G5 noticed the presence of inverted faces in a control no-report block at the end of the experiment, while G18 did not notice them. Note that G22, for whom stimulus intensity in experiment 3 could be decoded in the lateral visual cortex, also noticed the inverted faces. In the future, it will be interesting to cross-validate these results with oculometric measurements, which provide a way to classify whether a stimulus is perceived in no-report conditions (Frässle et al., 2014; Hesse & Tsao, 2020; Kronemer et al., 2022; White et al., 2022). It will also be important to inverse the order of report and no-report conditions to rule out that participants engaged in post-perceptual cognitive processing during experiment 3 after having performed the task extensively during experiment 1 (Block 2019).

### Slow accumulation for threshold stimuli produce late markers of conscious access in the VVC

In our detection tasks, the onset of the HGA detection effect in the VVC was consistent across experiments (0.33 s after stimulus onset in experiment 1 and 0.34 s in experiment 2) and reflected late reaction times (0.74±0.11 s, with only 2.16% of trials under 0.4 s). Late timings are consistent with what was observed in single neurons in the parietal cortex in a vibrotactile detection task using threshold stimuli (Pereira et al., 2021). Despite these late timings, our modeling results suggest that evidence accumulation could be starting as early as 150 ms after stimulus onset. Our findings thus raise the possibility that the neural activity underlying perceptual experience could occur at different latencies after the stimulus onset according to both the drift (deterministic) and the diffusion (stochastic) components of the evidence accumulation process. In experiment 1, we observed that the HGA increase in the VVC was markedly delayed for slower responses times (i.e., varied according to the diffusion component). In experiment 2, we did not have access to response times but we observed an interaction effect between detection responses and stimulus intensity, suggesting that the increase in HGA was delayed following threshold compared to suprathreshold stimuli. Accordingly, on the one hand, non-degraded face stimuli should have strong drift rates and lead to early increases of neural activity in the VVC (Axelrod et al., 2019; 2022; Broday-Dvir et al., 2023; Cogitate Consortium et al., 2023; Khuvis et al., 2021; Lachaux et al., 2005; Minxha et al., 2017; Quiroga et al., 2008; Quiroga et al., 2023; Schrouff et al., 2020, Vishne et al., 2023). On the other hand, in our study, degraded faces at the perceptual threshold are associated with low drift rates that delay neural activity correlating with conscious access. Our results therefore call for a nuanced interpretation of latencies, particularly as a diagnostic tool for assessing theories of consciousness (Cogitate Consortium, 2023).

### A leaky evidence accumulation process can account for the pattern of results observed in the VVC

It has been proposed that contents reach conscious access when an accumulated signal crosses a threshold (Dehaene, 2009; Moutard et al., 2015; Pereira et al., 2022; Shadlen & Kiani, 2011), and a growing body of evidence has linked evidence accumulation to perceptual experience (e.g., Kang et al., 2017; Nie et al., 2023; Pereira et al., 2021; Tagliabue et al., 2019). Our study adds to this body of work: The neural code that was found to reflect evidence accumulation in the VVC also reflected conscious access during passive viewing, consistent with fMRI (Dellert et al., 2021), intracranial (Broday-Dvir et al., 2023; Vishne et al., 2023) and monkey electrophysiology (Hesse & Tsao, 2020) studies showing activity in occipitotemporal regions in visual no-report paradigms. Also, the occurrence of false alarms was found to correlate with the number of times that the decoded latent variable crossed a detection threshold in the VVC (**Figure 5**). These results fit into a recent proposal that several aspects of perceptual experience are determined by a leaky evidence accumulation process (Pereira et al., 2022). To formally gauge whether it accounts for conscious access in our study, we assessed if simulations based on a computational model instantiating this process could reproduce the observed pattern of data. Despite being trained on behavioral data only, this model reproduced several idiosyncratic features of the HGA in the VVC (**Figure 7**). The results therefore provide support for the proposal that conscious access is determined by a leaky evidence accumulation instantiated in high-level sensory regions. At this stage, this process appears compatible with several of the main candidate theories of consciousness (Pereira et al., 2022), although it will be helpful to refine their experimental predictions to determine how exactly evidence accumulation fits into them (Seth & Bayne, 2022).

### Confidence computations in the VVC are related to the maximum of accumulated evidence

Some authors have argued that perceptual experience consists not only of conscious access, but also of distinct epistemic feelings of confidence associated with perceptual monitoring (Dokic & Martin, 2015; Pereira et al., 2022). Our results show that the same neural code reflects conscious access and perceptual confidence. Perceptual confidence was found to relate to the maximal level of evidence that is reached after the threshold is crossed when participants see a face. Consistent with the view that confidence derives from maximal evidence (Pereira et al., 2021; 2022), decoding of confidence in the VVC was best around the time of the maximum of the latent variable (**Figure 4E**), a direct indication that this information was important in confidence computations. Our data indicated that the SPC and IFC also reflected perceptual confidence, but hinted at different computations across regions. Contrary to what we observed in the VVC, we found that confidence was best predicted early in the trial in the SPC. In the IFC, a region that has been shown to accumulate evidence for confidence (Goueytes et al., 2024; Noppeney et al., 2010), confidence could not be decoded at the time of the maximum of the latent variable. These results indicate that confidence is represented differently in those regions compared to what is observed in the VVC and what is predicted by the leaky evidence accumulation process. An interesting avenue for future research will be to disentangle the roles of those different regions and their interactions in how confidence computations unfold.

## Conclusion

We showed that a neural code in the ventral visual cortex reflected evidence accumulation for face perception, also when reports were delayed or absent, confirmed by a computational model based on a leaky accumulation process that could reproduce traces of high gamma activity in all three experiments. Additionally, it could decode confidence judgments in seeing a face. Altogether, our results show that evidence accumulation occurs beyond decision-making and could explain the perceptual experience associated with a stimulus. This proposal also implies that conscious access might occur at varying latencies after stimulus onset, depending both on its physical intensity and the stochastic nature of the associated sensory evidence.

## Acknowledgments

The authors would like to thank Clarissa Baratin, Blandine Chanteloup, Manik Bhattacharjee, Martin Kojan, Pavel Daniel and Jan Cimbálník for their help with the electrode localization procedure, Emmanuelle Marmet, Marine Carmona, and Fosco Bernasconi for their help with data collection, and Megan Peters and Colin Hoy for useful feedback on draft versions of this article. DG is supported by a Marie Skłodowska-Curie Action funded by the European Commission through Horizon Europe (MetaChange, project 101110783 - HORIZON-MSCA-2022-PF-01-01). MB is supported by project nr. LX22NPO5107 (MEYS): Funded by European Union-Next Generation EU and Czech Science Foundation, project 22-28784S. MP was supported by a Postdoc.Mobility fellowship from the Swiss National Science Foundation (P400PM_199251). NF has received funding from the European Research Council (ERC) under the European Union’s Horizon 2020 research and innovation programme (Grant agreement No. 803122).

## Authors’ contributions

Conceptualization and methodology: FS, MP and NF; software and code validation: MP.; investigation: FS, LJ, DG, MR, RM and RR.; formal analysis: FS, RM and MP; resources: MB, LMi, PK and NF; supervision: LMu, NF and MP; funding acquisition: NF; visualization: FS; writing - original draft: FS, MP, NF; writing - reviewing and editing: all authors.

## Methods

### Resource availability

Data and code availability: All data and code are available on the Open Science Framework (https://doi.org/10.17605/OSF.IO/8QE3A).

Lead contact: Further requests for resources should be directed and will be fulfilled by the lead contact, François Stockart.

### Participants and data acquisition

29 participants (10 females, aged 36±9 years; **Supplementary Table 1**) with drug-resistant focal epilepsy were included across two separate sites, the Grenoble Alpes University Hospital, France (n = 22) and St. Anne’s University Hospital Brno, Czech Republic (n = 7). They were implanted with 10 to 18 stereotactic electrodes in the context of presurgical evaluation, and electrode placement was conditioned on clinical criteria. Electrode implantation and participation in research protocols received ethical approval from CPP du Sud-Ouest et Outre-Mer 4 CPP18-001b/2017-A03248-45 (in Grenoble) and Ethical committee at St. Anne’s University Hospital 46V/2019 (in Brno). Participants were implanted with semirigid, multilead electrodes of 0.8 mm diameter, containing up to 18 contact leads 2-mm-wide and 1.5-mm-apart (DIXI Medical® USA). Neural recording in Grenoble was conducted at 512 Hz with the Micromed (Trevisio, Italy) audio-visio-EEG recording system. Neural recording in Brno was conducted at 5000 Hz with the M&I (BrainScope, Prague, Czech Republic) 192-channel research EEG acquisition system, and downsampled to 512 Hz before preprocessing. The anatomical locations of electrode contacts in both sites were identified on BrainVisa (Geffroy et al., 2011) using a post-implantation CT scan co-registered with a pre-implantation T1 MRI scan.

Four participants were excluded from all analyses because they did not respond directly after stimulus presentation in experiment 1 and/or had a higher rate of false alarms than hits in experiment 2. For two of the included participants (G2 and G8) we could not acquire electrophysiological recordings in experiment 3 due to clinical constraints. Finally, one participant (G5) came back to the hospital with a new implantation scheme and was tested again on this occasion. We report the results for the two sessions from this participant as two independent datasets.

### Stimuli

Stimuli were presented on a Dell Precision 7550 laptop (15 inches), with 1920 x 1080 pixels resolution and 60 Hz refresh rate in Grenoble, and on a 27 inches LED display with 1920 x 1080 pixels resolution and 120 Hz refresh rate in Brno. We checked that the behavioral results were similar in the two data collection centers despite the different equipment used by adding the recording center as a variable of no-interest in a control analysis. The fixation cross was presented at the center of the screen, and each of its four arms was 10 pixels long and 2 pixels wide. Mask frames were also centered on the screen and measured 768 x 768 pixels. For each trial, the masks were generated using one of four greyscale pictures of neutral-valence human faces (2 males, 2 females) that were cropped such that they contained only facial features and blurred around the eyes and mouth. Scrambled images were obtained by randomly permuting the phase angle of the discrete Fourier transform of the images, on which an inverse discrete Fourier transform was applied. The phase scrambled stimulus generated for each frame was combined with that of the original face image at an intensity equal to I, using I*face + (1-I)*mask.

Face stimuli in target-present trials in experiment 1 were all presented at detection threshold (threshold trials) for three frames (600 ms). In experiments 2 and 3 faces could be presented at threshold or above threshold (suprathreshold trials). Intensity I in suprathreshold trials was 1.25 and 1.5 times the threshold value in experiment 2 and 3, respectively. Detection threshold was determined using a 1-up/1-down staircase procedure titrating 50% hit rate, with an initial I value of 0.15. Stimulus intensity was increased/decreased by a percentage of its current value (5%) after each hit/miss response in target-present trial. In order to account for possible threshold drifts during the experiment, the I value was also continuously adapted in experiment 2 after each threshold trial by an increase after “seen” and a decrease after “not seen” responses. The step in experiment 2 was 5% for participants B1 to B7 and G1 to G8, and decreased to 2% for participants G9 to G21 to improve staircase convergence.

In target-absent trials, the I value was set at 0 for all mask frames. In experiment 2, there were some trials (one per block) for which a second face stimulus was presented, also for three consecutive frames. In those trials, the two faces were always presented at suprathreshold value (i.e., 1.25 times the threshold value), and also always had the same onset relative to the beginning of the sequence, at 200 and 1400 ms.

### Procedure

Participants were seated in a hospital bed, with the stimulation laptop in front of them on a bedside table. The experiments were run with Psychtoolbox-3 (Brainard, 1997) on Matlab 2019b. The stimulus sequence was the same in all three experiments. First, participants were presented with the fixation cross on a gray screen background for 600 ms. Then, 13 consecutive mask frames were presented at 5 Hz for 2600 ms. A face stimulus was embedded for a duration of 600 ms starting at a random frame from 600 to 1800 ms after the onset of the sequence. This sequence was followed by a random delay of 500 to 1000 ms during which no stimulus was presented. In experiment 1, participants were asked to press a key as soon as they saw a face. In experiment 2, participants provided detection and confidence responses following a prompt which remained on screen until a response was made. The second answer screen was presented 100 ms after the first response, and the next trial started 100 ms after the confidence report. In experiment 3, participants passively observed the stimuli.

Before taking part in the three experiments, participants were presented with a five-trials long demonstration with the same structure as experiment 2. In the first trial, the embedded face stimulus was presented at maximum intensity (i.e., no phase scrambling at all). In subsequent trials, the intensity was reduced by steps of 20% so that they understood the difficulty of the task. Finally, in the last trial, the target was absent. Participants were instructed to wait for the answer screens to provide detection and confidence reports. If the instructions were still unclear to the participant after this short sequence, the demonstration was repeated once again before starting experiment 1.

#### Experiment 1

Participants were asked to provide an immediate response as soon as they saw a face and not to respond if they did not see a face. They responded by pressing the button “2” on a numerical keyboard. No feedback was given on their performance and they directly proceeded to the next trial. Participants performed two blocks of 52 trials in Grenoble and 40 trials in Brno, including ∼20% target-absent trials (10 and 8 trials in Grenoble and Brno, respectively).

#### Experiment 2

Participants were instructed to respond at the end of the trial, when a question displayed in the middle of the first answer screen probed them for a detection report (“How many faces did you see?”). They responded by pressing one of three buttons on a numerical keyboard corresponding to “none”, “one”, or “more than one”. They were informed that in some trials two faces may be presented. These trials occurred only once per block. Only “one face” responses were treated as hits or false alarms when a single face was presented during the trial. A second answer screen then asked participants to report their confidence on a three-point scale conditioned to the detection report (“Confidence of having seen no face” or “Confidence of having seen one face or more”). They responded by pressing another set of three buttons corresponding to “unsure”, “moderately sure”, or “very sure”. Participants were encouraged to make full use of the confidence scale, both when they had reported seeing a face and when reporting seeing no face. When they reported seeing “more than one face” to the detection question, they were asked to report their confidence based on the clearest face stimulus they had perceived. In Grenoble, participants performed three blocks. In Brno, they performed two blocks. Each block consisted of 59 trials, including 10 target-absent trials and one trial with two faces. Target-present trials were constituted of an equal number of threshold and suprathreshold trials.

#### Experiment 3

The stimulus sequence was the same as in the previous two experiments, yet here, participants were not asked to report anything. Instead, they were simply instructed to keep fixation and remain attentive to the stimulation. There were no target-absent trials. Akin to experiment 2, there was an equal number of threshold and suprathreshold trials. Given that no answers were provided, stimulus intensity was not titrated. The number of blocks was identical to Experiment 2, with 48 trials in each block. To test if participants were attentive to the visual stream, they were presented with a short no-report control block at the end of experiment 3. The block consisted of 12 trials, with 8 upright and 4 inverted face stimuli (equal number of threshold and suprathreshold trials). We assumed that if they saw them, the inverted faces would be surprising to the participants, as all other faces presented to them previously were upright. At the end of the block, the experimenter asked the participant if they noticed anything odd about the last block. If they responded that it was the case, they were further asked what that was. A record was kept of whether they reported seeing inverted faces or not.

### Behavioral analysis

Trials with early or late responses in experiment 1 were excluded from all analyses. A response was considered early if it occurred earlier than 400 ms prior to stimulus onset and as late if it occurred later than 1500 ms after stimulus onset. On average, these responses accounted for 2.16±2.32% and 0.86±2.03% of trials, respectively, and these trials were not included in the analyses. In experiment 2, trials with response times over 10 seconds were excluded (0.54% of trials), as it was likely that the participant was not performing the task anymore if they took so long to respond. We also excluded trials where more than one face was presented and trials where participants reported seeing more than one face (6.87% of remaining trials where one face was presented).

Detection and confidence responses in experiment 2 were fitted with effects-coded generalized linear mixed effects regressions with an underlying binomial distribution and a logit link function (‘fitglme’ function in Matlab). For detection responses, the relative intensity of the stimulus (threshold vs. suprathreshold) was specified as fixed effect factors and participants as random effects. For decision confidence responses, the fixed effects were relative intensity, detection response and their interaction. Separate models were then fit for hit and miss trials. We found that most participants reported a majority of “very sure” responses (**Supplementary figure 5**). To avoid design cells containing very few trials, we split the scale in two, reporting “very sure” responses as “high confidence” and pooling “moderately sure” and “unsure” responses as “low confidence”. Log likelihoods were compared to determine whether to add random coefficients to each model.

### Electrophysiological data processing and analysis

#### Preprocessing

The location of contacts was determined using a pre-implantation MRI and a post-implantation CT scan on the Intranat software (Deman et al., 2018). MarsAtlas parcellations were performed in participants’ native brain space (Auzias et al., 2016). Coregistration to MNI space was performed in Intranat using SPM12.

Electrophysiological signals were visualized and analyzed under Matlab 2020b using custom-made scripts and the FieldTrip toolbox (Oostenveld et al., 2011). Raw signals were visually inspected to exclude channels with artifactual or epileptic activity (24.03% of all contacts). A notch filter was then applied at 50 Hz, 100 Hz and 150 Hz to eliminate power line artifacts. Bipolar derivations were obtained by re-referencing neighboring contacts to each other. When one isolated contact was excluded, we computed a derivation for its two neighboring contacts. Of 5262 recording contacts, 3572 channels remained after inspection and bipolar derivation. Channels that were not located in the MarsAtlas (n = 271) were excluded from all analyses.

High-gamma activity (HGA) was computed based on the electrophysiological signal between 70 and 150 Hz. Non-causal filters were applied on each trial to define 20 Hz overlapping frequency sub-bands (70 to 90 Hz, 80 to 100 Hz, etc.), whose envelope was extracted with a Hilbert transformation. The extracted signal was then divided by the mean activity starting one second before the trial and ending at trial onset, and averaged across sub-bands. The resulting signal was smoothed using a quadratic Savitzky-Golay filter (Savitzky & Golay, 1964) of polynomial order 2 with a 200 ms window to preserve the peak amplitudes. Note that dividing with baseline activity means that the signal is centered around 1. Finally, the data were resampled at 64 Hz. Individual trials with remaining artifacts were removed by visual inspection (7.21% of trials).

#### Anatomical definitions

We limited our principal analyses to four pre-registered ROIs, which have been found to accumulate evidence (Kim & Shadlen, 1999; Pedersen et al., 2015; Ploran et al., 2007; Roitman & Shadlen, 2002; Tremel & Wheeler, 2015) and are suspected to play a role in perceptual experience (Pereira et al., 2021; Schrouff et al., 2020; van Vugt et al., 2018; Weilnhammer et al., 2021; Woolnough et al., 2020): the VVC, the SPC, the IFC and the dlPFC. ROIs were defined based on combinations of regions from the MarsAtlas cortical parcellation model (Auzias et al., 2016). The VVC (234 channels) corresponded to the caudal medial visual cortex, medial inferior temporal cortex and rostral inferior temporal cortex in MarsAtlas. The SPC (72 channels) corresponded to the superior parietal cortex and medial superior parietal cortex, the IFC (267 channels) to the rostral ventral premotor cortex and rostral ventrolateral prefrontal cortex, and the dlPFC (103 channels) to the rostral dorsolateral inferior prefrontal cortex and rostral dorsolateral superior prefrontal cortex. Responsive channels in ROIs were visualized in native space by a trained neurologist (A. R.). To make sure of the anatomical origin of the observed effects, we performed two sets of control analyzes. First, because some channels classified as belonging to the IFC were actually located in the anterior insula, we looked at a control region encompassing the anterior insular cortex and the functionally homologous internal operculum (Evrard, 2020). Second, we looked at the impact of excluding channels that were located in the white matter. We also performed exploratory analyses on all regions in the MarsAtlas that included more than 30 channels and at least 10 responsive channels in the hit vs. miss contrast in experiment 1 (21 out of 31 possible regions). Plots of brain slices were created using SPM12 (Ashburner, 2009).

#### Univariate analyses

We performed nonparametric clustering analyses on individual channels for hit vs. miss trials or threshold vs. suprathreshold trials (Maris & Oostenveld, 2007). Responsive channels (α = 0.05) were identified as those showing a positive cluster (i.e., HGA in hit trials > HGA in miss trials) in experiment 1. In the evidence accumulation analysis, HGA slopes were calculated by fitting a linear regression model starting 200 ms after stimulus onset and stopping 100 ms before the response. HGA slopes were then fitted to reaction times using linear mixed-effects regressions.

For analyses based on ROIs, in experiments 2 and 3, the HGA of each responsive channel was averaged across time points for each condition within the positive cluster identified for that channel. This averaged HGA was then fitted with generalized linear models with an underlying gamma distribution and a log link function. Time-resolved results were obtained by fitting a model on each time point in a one-second window following stimulus presentation and correcting across time points using the false discovery rate procedure (Benjamini & Hochberg, 1995). In experiment 1, HGA slopes were fitted with hierarchical mixed-effects regressions, considering that slopes followed a normal distribution under the null hypothesis. To account for the full hierarchical structure of the data, all analyses on ROIs involved hierarchical models with random effects for channels and participants. As an example, the model using detection and stimulus intensity as predictors in experiment 2 had the following structure:

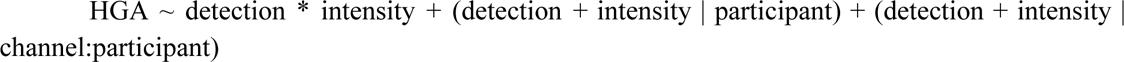

#### Multivariate analyses

Custom decoders were trained to discriminate between hit and miss trials in experiment 1 separately for each participant, on ROIs with five channels or more. They were fed trial-by-trial HGA from a window starting at stimulus onset and finishing 1 s later. The Nelder-Mead method was used to obtain weights that maximized the distance of the maximum values of a time-resolved trial-by-trial latent variable across classes. Specifically, we obtained a distance term for each trial by calculating the logarithm of the normalized distance between the maximum and a threshold. To counteract the effect of class imbalances, the distance term of a given trial was divided by the number of trials in that trial’s class. The error term the fit function attempted to maximize consisted of the difference between distance terms in class 1 (e.g. hit) and class 2 (e.g., miss). A penalty was applied on the overall error term to promote sparsity (‘L1 regularization’). A Bayesian optimization search was performed to find the best value of that hyperparameter, using a 5-fold cross-validation procedure.

Decoders performing within-sample predictions were trained using a leave-one-out procedure. Other decoders were trained on all trials in experiment 1 to perform out-of-sample predictions on detection response, confidence judgements and stimulus intensity in experiments 2 and 3. In the case of confidence judgment and stimulus intensity, the labels were modified such that a successful prediction corresponded to classifying a high confidence/suprathreshold trial as a hit or a low confidence/threshold trial as a miss. Decoders were only tested when there were at least 5 trials per experimental cell (e.g., low and high confidence). Decoders were tested in the same stimulus-locked window than the one they were trained on (0 to 1000 ms after face stimulus onset), except in analyses including trials without such events to lock the window on, i.e., false alarms or reports of seeing more than one face, in which case the decoders were tested on a window starting at the earliest time that faces could appear and finishing right before the earliest possible onset answer screen (600 ms to 2800 ms following the start of the trial). In confidence analyses, we also tested decoders’ performance in a window of 400 ms centered on the time of the maximum of the latent variable. To assess decoding performance, we reported the area under the receiving operator characteristics (AUROC). We permuted labels 1000 times to assess whether the obtained values for each participant were significant. To correct for multiple comparisons within each region of interest, we performed the false discovery rate procedure across participants (Benjamini & Hochberg, 1995). For group analyses, we tested observed average AUROC values against a null distribution obtained by averaging AUROC values obtained from permutations. To correct for multiple comparisons, we again applied the false discovery rate, yet this time across regions (Benjamini & Hochberg, 1995).

We estimated a latent variable for each trial by applying the set of weights from the decoder at each point in time. The threshold obtained in the training procedure did not provide the best categorization of classes, as it was focused on maximizing distances instead. We thus estimated individualized detection thresholds. per participant by taking the threshold value of the receiver operating characteristic curve that maximized the rate of true positives (hit trials correctly classified as such) to false positives (miss trials incorrectly classified as hit). In evidence accumulation analyses, hit trials for which the estimated latent variable crossed the threshold were selected for a Spearman’s rank-order correlation between the timing of this threshold crossing and reaction times. In analyses on false alarms and trials in which participants reported seeing more than one face (**Figure 5**), we computed and averaged by condition the number of times that the latent variable crossed the detection threshold computed from hit and miss trials in experiment 2. Statistical significance was also calculated using permutation tests.

### Computational model

The model instantiated an evidence accumulation process with a leakage factor (Usher & McClelland, 2001; Pereira et al., 2021; 2022) and included five free parameters: a drift rate with mean Ɣ and standard deviation *s*, a flat decision boundary *θ*, a leakage factor λ, and a non-decision time *ndt*. A white Gaussian noise process *W*(*t*) with zero mean and unit variance was added to the process to account for stochasticity (Eq.1). Since neuronal responses such as high-gamma activity are positive, the evidence accumulation EA(t) process was bound to zero for biological plausibility.

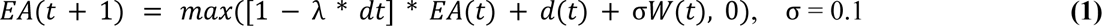

The accumulation process started at the beginning of each trial with a zero drift rate. On stimulus onset *S*, following a non-decision time, *d*(*t*) increased to a level (Ɣ + *sZ*)*log*(*I* + 1), for 600 ms (stimulus duration) before returning to zero at stimulus offset (Eq.2). *I* represents the stimulus intensity and Z a zero-mean and unit-variance Gaussian noise varying on a trial-by-trial basis.

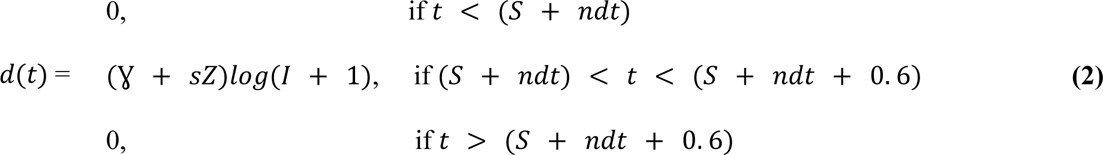

Stimuli were considered as detected if the accumulated evidence reached the decision boundary *θ.* Parameters were adjusted based on pilot data to reproduce behavioral and neural data (Ɣ = 0. 012; *s* = 0. 4 * Ɣ; θ = 3. 45; λ = 4; *ndt* = 0. 3).

We simulated 10000 trials including 4000 threshold trials, 4000 suprathreshold trials, and 2000 catch trials (no stimuli). As in the actual experiment, suprathreshold intensity was set to 1.25 times the threshold intensity for experiment 2 and 1.5 times that intensity for experiment 3. The non-decision time was also divided into two components: a pre-decision time of 150 ms and a post-non-decision time of 150 ms. The set of parameters was used to generate traces of accumulating evidence (time-locked to stimulus onset) and simulate detection responses. A fixed random time shift (normal distribution with a mean of 0 and a standard deviation of 50 ms) was applied to each trial’s evidence accumulation trace to simulate non-decision time variability, similar to previous works (Van den Berg et al., 2016). Similar to behavioral analysis, simulated response times outside the range of 0.4 s to 1.5 s following stimulus onset (7.6% and 5.4%, respectively) were excluded.

For visualization, we separated simulated hit trial traces (for threshold condition) in three terciles based on reaction times to show the relationship between response times and accumulating evidence in experiment 1. We plotted the average of accumulating evidence in each tercile from 0.25 s before stimulus onset until the median reaction time (**Figure 7B** left panel).

## Supplementary tables

**Supplementary table 1.** Demographic information about participants. The letter in the participant’s ID refers to the center: “G” for Grenoble and “B” for Brno. Antiseizure medication corresponds to the daily dose prescription.

| ID | Sex | Age (years) | Hand laterality | Antiseizure medication |
| --- | --- | --- | --- | --- |
| G1 | M | 35 | R | LACOSAMIDE 600mg |
| G2 | M | 32 | R | PERAMPANEL 6mg; CARBAMAZEPINE 1000mg |
| G3 | M | 25 | R | LAMOTRIGINE 500mg; ZONISAMIDE 100mg; ZEBINIX 800mg |
| G4 | M | 26 | R | OXCARBAZEPINE 1800mg; PERAMPANEL 8mg |
| G5 | F | 25 | L | PERAMPANEL 6mg; LACOSAMIDE 400mg; LAMOTRIGINE 175mg |
| G6 | M | 24 | L | ZONISAMIDE 200mg; CARBAMAZEPINE LR 800mg; 1600mg; LEVETIRACETAM 1750mg; CLONAZEPAM 1.5mg |
| G7 | M | 38 | R | LAMOTRIGINE 600mg; LACOSAMIDE 400mg |
| G8 | F | 52 | R | CARBAMAZEPINE LR 800mg |
| G9 | F | 38 | R | LAMOTRIGINE 400mg; CARBAMAZEPINE LR 1000mg |
| G10 | M | 56 | R | PARAMPANEL 4mg; ZONISAMIDE 400mg |
| G11 | M | 39 | R | ESLICARBAZEPINE 1200mg; PERAMPANEL 12mg |
| G12 | M | 41 | R | CARBAMAZEPINE LR 800mg; LAMOTRIGINE 300mg; BRIVARACETAM 25mg |
| G13 | F | 48 | R | LACOSAMIDE 400mg; LAMOTRIGINE 300mg |
| G14 | M | 28 | R | ZONISAMIDE 400mg; ESLICARBAZEPINE 120mg |
| G15 | M | 39 | R | LACOSAMIDE 600mg; PERAMPANEL 6mg |
| G16 | M | 37 | R | CARBAMAZEPINE LR 1200mg |
| G17 | F | 45 | L | CARBAMAZEPINE LR 1200mg; LEVETIRACETAM 3000mg; CLONAZEPAM 1mg; TOPIRAMATE 300mg |
| G18 | M | 27 | R | LEVETIRACETAM 2000mg; CARBAMAZEPINE LR 800mg |
| G19 | F | 24 | L | LAMOTRIGINE 400mg |
| G20 | F | 36 | R | BRIVARACETAM 200mg |
| G21 | F | 48 | R | LAMOTRIGINE 300mg; CENOBAMATE 100mg |
| G22 | M | 33 | R | CARBAMAZEPINE LR 400mg; PREGABALINE 400mg; TOPIRAMATE 300mg |
| B1 | M | 38 | R | LAMOTRIGINE 400mg; LEVETIRACETAM 2000mg |
| B2 | M | 39 | L | LAMOTRIGINE 400mg; ZONISAMIDE 250mg |
| B3 | M | 30 | R | LACOSAMIDE 500mg; BRIVARACETAM 400mg; PERAMPANEL 10mg; ESLICARBAZEPINE 800 mg |
| B4 | F | 26 | L | BRIVARACETAM 200mg; PERAMPANEL 10mg |
| B5 | F | 31 | R | BRIVARACETAM 200mg; ESLICARBAZEPINE 800mg; PERAMPANEL 6mg |
| B6 | M | 30 | R | LACOSAMIDE 400mg; BRIVARACETAM 200mg; PERAMPANEL 4mg |
| B7 | M | 41 | R | CARBAMAZEPINE LR 900mg; PERAMPANEL 4mg |

**Supplementary table 2.** The number of included channels corresponds to contact sites within regions of interest after signal inspection, bipolar derivation and exclusion of contacts that were not localized in Mars Atlas.

| ID | N° of recording sites | N° of included channels | N° of channels in VVC | N° of channels in SPC | N° of channels in IFC | N° of channels in dIPFC |
| --- | --- | --- | --- | --- | --- | --- |
| G1 | 122 | 101 | 15 | 0 | 0 | 0 |
| G2 | 220 | 173 | 29 | 8 | 0 | 0 |
| G3 | 235 | 190 | 14 | 6 | 6 | 0 |
| G4 | 201 | 134 | 4 | 0 | 14 | 11 |
| G5 | 207 | 177 | 16 | 0 | 27 | 6 |
|  | 221 | 175 | 0 | 0 | 31 | 0 |
| G6 | 231 | 183 | 10 | 36 | 12 | 0 |
| G7 | 192 | 112 | 1 | 0 | 5 | 3 |
| G8 | 176 | 139 | 6 | 12 | 7 | 0 |
| G9 | 161 | 91 | 0 | 0 | 18 | 0 |
| G10 | 141 | 111 | 14 | 0 | 10 | 8 |
| G11 | 200 | 124 | 0 | 0 | 17 | 8 |
| G12 | 202 | 127 | 16 | 0 | 17 | 0 |
| G13 | 169 | 62 | 2 | 1 | 7 | 7 |
| G14 | 197 | 102 | 0 | 5 | 14 | 0 |
| G15 | 143 | 38 | 0 | 1 | 4 | 3 |
| G16 | 156 | 83 | 0 | 0 | 15 | 4 |
| G17 | 132 | 73 | 14 | 0 | 1 | 0 |
| G18 | 185 | 108 | 19 | 0 | 14 | 10 |
| G19 | 186 | 80 | 9 | 0 | 0 | 0 |
| G20 | 186 | 111 | 9 | 3 | 3 | 4 |
| G21 | 158 | 108 | 3 | 0 | 8 | 3 |
| G22 | 176 | 87 | 11 | 0 | 0 | 0 |
| B1 | 168 | 124 | 24 | 0 | 0 | 14 |
| B2 | 118 | 34 | 7 | 0 | 0 | 0 |
| B3 | 165 | 122 | 0 | 0 | 17 | 9 |
| B4 | 172 | 94 | 9 | 0 | 1 | 1 |
| B5 | 172 | 125 | 5 | 0 | 11 | 0 |
| B6 | 137 | 37 | 8 | 0 | 0 | 3 |
| B7 | 133 | 76 | 0 | 0 | 8 | 9 |

**Supplementary table 3.** Decoding performance (AUROC) for detection in experiment 1, evidence accumulation as assessed by the correlation (Spearman’s rho) between reaction times and the timing at which the latent variable crosses a threshold, decoding performance for detection in experiment 2, decoding performance for intensity in experiment 3. This table only shows participants with significant decoding performance in experiment 1 in the four ROIs. Missing values (n/a) are caused by some participants not being able to participate in experiment 3 due to clinical constraints. P-values were obtained with permutation tests and corrected with false discovery rate across participants in the same region: *** = p < 0.001, ** = p < 0.01, * = p < 0.05.

| Region | ID | Decoding of detection in exp 1 (AUROC) | Evidence accumulation in exp 1 (Spearman's rho) | Decoding of detection in exp 2 (AUROC) | Decoding of intensity in exp 3 (AUROC) |
| --- | --- | --- | --- | --- | --- |
| VVC | G2 | 0.8*** | 0.64* | 0.71*** | n/a |
|  | G5 (ses 1) | 0.81*** | 0.33* | 0.77*** | 0.69** |
|  | G8 | 0.83*** | 0.3* | 0.86*** | n/a |
|  | G18 | 0.85*** | 0.46* | 0.77*** | 0.57 |
| SPC | G2 | 0.75** | 0.49 | 0.71*** | n/a |
|  | G3 | 0.69*** | -0.16 | 0.58 | 0.56 |
|  | G6 | 0.61* | 0.3* | 0.58 | 0.55 |
|  | G8 | 0.81*** | 0.41* | 0.8*** | n/a |
|  | G14 | 0.63* | -0.22 | 0.64** | 0.5 |
| IFC | G3 | 0.86*** | 0.26 | 0.6 | 0.49 |
|  | G4 | 0.66* | 0.35* | 0.49 | 0.58 |
|  | G5 (ses 1) | 0.75*** | 0.47* | 0.56 | 0.56 |
|  | G5 (ses 2) | 0.72** | 0.19 | 0.6 | 0.57 |
|  | G8 | 0.82*** | 0.45* | 0.77*** | n/a |
|  | G9 | 0.86*** | 0.46* | 0.65* | 0.6 |
|  | G10 | 0.73*** | 0.38 | 0.64* | 0.51 |
|  | G12 | 0.71*** | 0.19 | 0.57 | 0.5 |
|  | G13 | 0.84*** | 0.4* | 0.54 | 0.43 |
|  | G16 | 0.85*** | 0.44* | 0.78*** | 0.55 |
|  | G18 | 0.84*** | 0.11 | 0.5 | 0.5 |
|  | G21 | 0.62* | -0.07 | 0.49 | 0.48 |
|  | B5 | 0.85*** | 0.43* | 0.6 | 0.52 |
| DLPFC | G10 | 0.74*** | 0.13 | 0.69*** | 0.55 |
|  | G13 | 0.69** | 0.54*** | 0.64* | 0.53 |
|  | B1 | 0.8** | -0.32 | 0.54 | 0.47 |
|  | B3 | 0.67*** | -0.03 | 0.65** | 0.58 |

**Supplementary table 4.** Same results as in *Supplementary table 3*, but for the lateral visual cortex (VCl)

| Region | ID | Decoding of detection in exp 1 (AUROC) | Evidence accumulation in exp 1 (Spearman's rho) | Decoding of detection in exp 2 (AUROC) | Decoding of intensity in exp 3 (AUROC) |
| --- | --- | --- | --- | --- | --- |
| VCI | G2 | 0.82** | -0.23 | 0.68** | n/a |
|  | G3 | 0.65** | 0.5* | 0.65*** | 0.48 |
|  | G6 | 0.63* | 0.16 | 0.57 | 0.49 |
|  | G8 | 0.76*** | 0.47* | 0.81*** | n/a |
|  | G19 | 0.72*** | 0.15 | 0.54 | 0.59 |
|  | G22 | 0.71*** | 0.24 | 0.77*** | 0.64** |

## Supplementary figures

**Supplementary figure 1.**
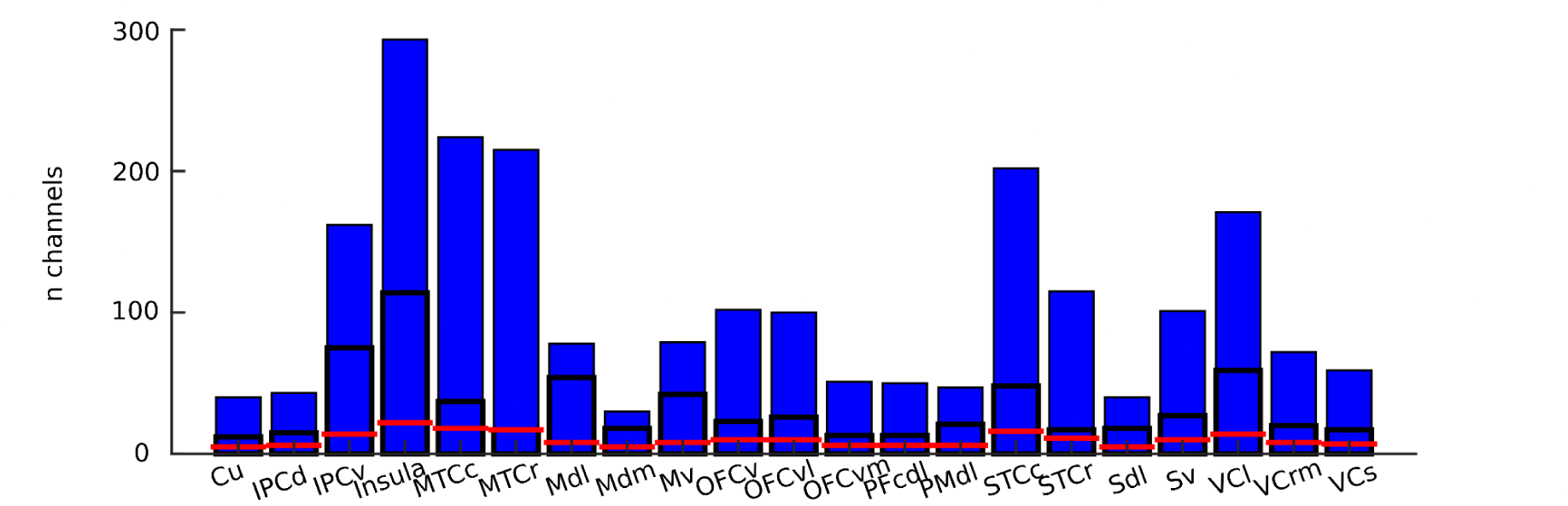
The total number of channels in other brain areas than the four regions of interest. Only areas with more than 30 channels overall and more than 10 responsive channels are included. The box with black contours shows the proportion of responsive channels, and the red line indicates the maximum number of responsive channels that would be expected by chance (α = 0.05). Names refer to: Cuneus (Cu), dorsal inferior parietal cortex (IPCd), dorsal inferior parietal cortex (IPCd), ventral inferior parietal cortex (IPCv), Insula, caudal middle temporal cortex (MTCc), rostral middle temporal cortex (MTCr), dorsolateral motor cortex (Mdl), dorsomedial motor cortex (Mdm), ventral motor cortex (Mv), ventral orbitofrontal cortex (OFCv), ventrolateral orbitofrontal cortex (OFCvl), ventromedial orbitofrontal cortex (OFCvm), caudal dorsolateral prefrontal cortex (PFcdl), dorsolateral premotor cortex (PMdl), caudal superior temporal cortex (STCc), rostral superior temporal cortex (STCr), dorsolateral somatosensory cortex (Sdl), ventral somatosensory cortex (Sv), lateral visual cortex (VCl), rostral medial visual cortex (VCrm) and superior visual cortex (VCs).

**Supplementary figure 2.**
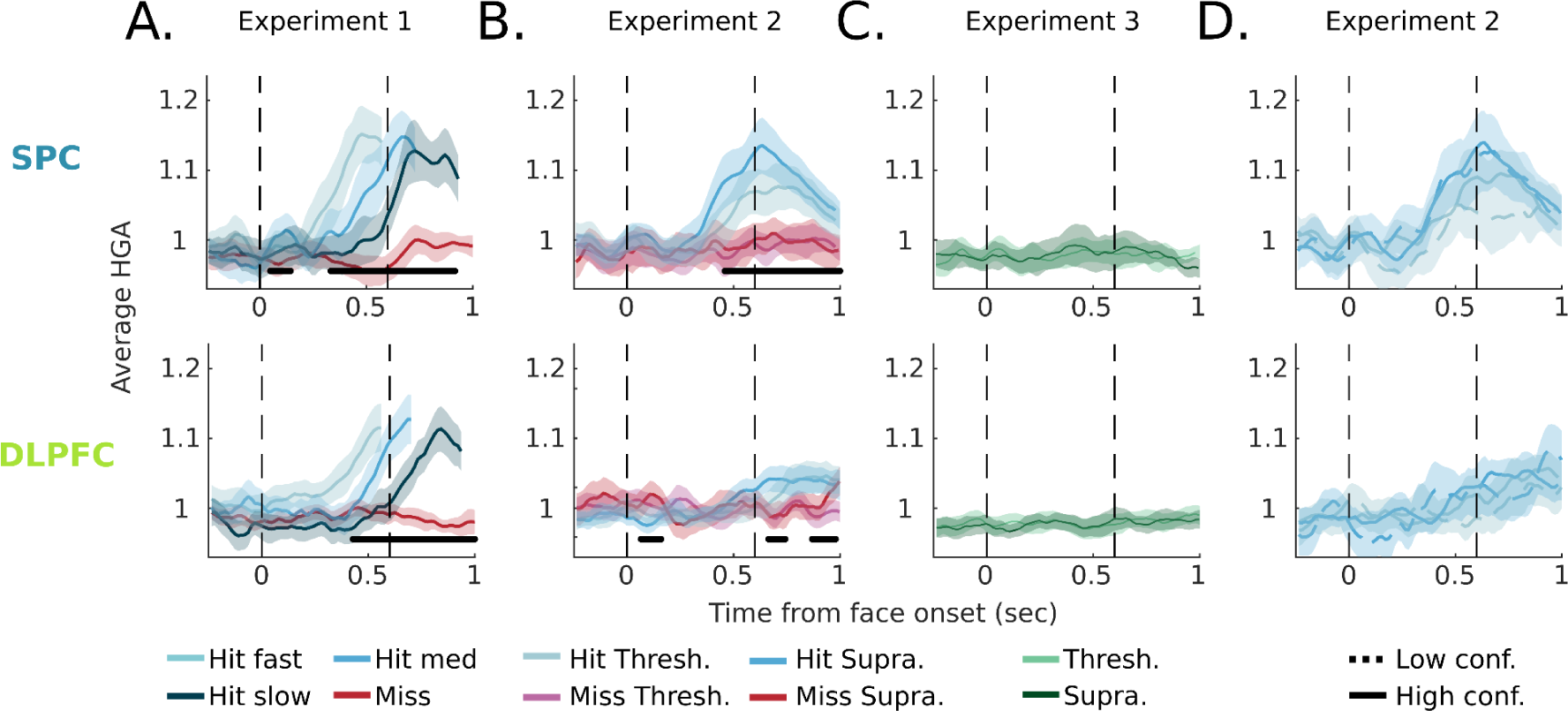
Time-resolved ROI results in SPC (top panels) and DLPFC (bottom panels) in the three experiments. *A.* The averaged activity in miss trials (red traces) and in hit trials in experiment 1, with hit trials separated in three terciles based on reaction times (blue traces) across all responsive channels. The black bars indicate significant main effects of detection as assessed with sample-by-sample hierarchical mixed-effects regressions (corrected for false discovery rate across time). *B.* The averaged activity in experiment 2, separated by first-order response type (hit vs miss) and intensity (threshold vs. suprathreshold), across all responsive channels. The black bars indicate significant main effects of detection. *C.* The averaged activity in experiment 3, separated by intensity (threshold vs. suprathreshold), across all responsive channels. *D.* The averaged activity in experiment 2 for low confidence and high confidence hit trials, across all responsive channels. All shaded areas indicate 95% confidence intervals.

**Supplementary figure 3.**
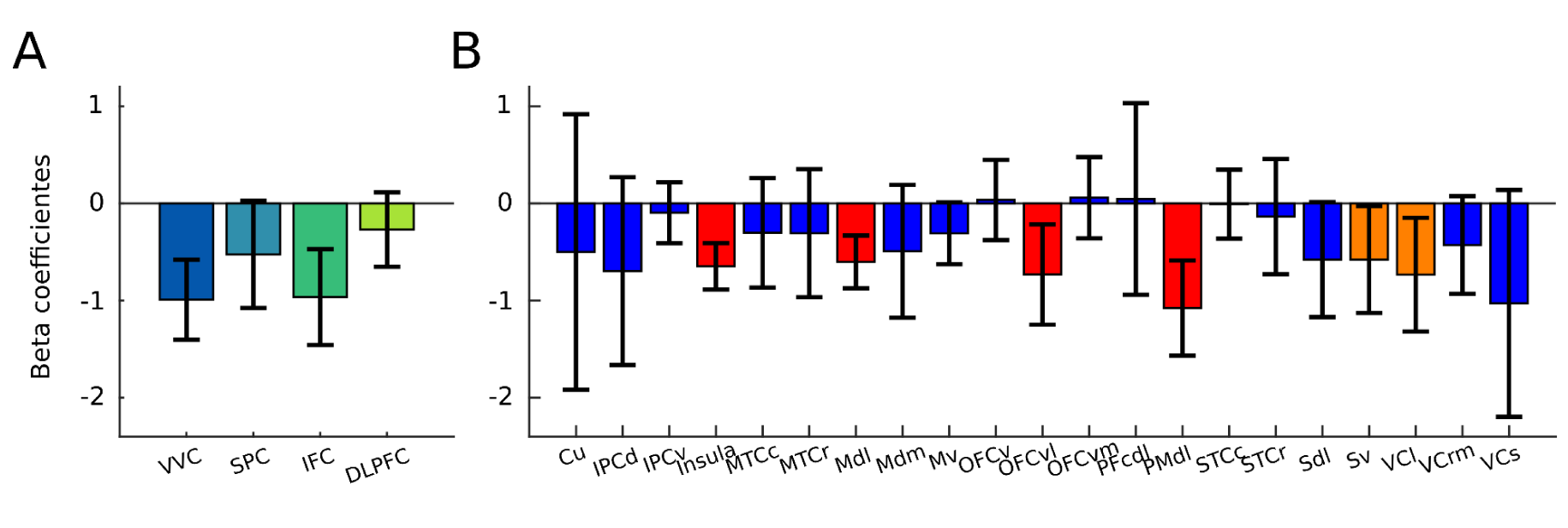
*A.* Beta coefficients for the analysis of HGA slopes as a function of reaction times analysis in the four ROIs in experiment 1. Error bars represent 95% intervals. *B.* Same plot as *A* for other brain areas. Blue boxes indicate regions where the correlation is non-significant, orange boxes regions where it is significant (uncorrected) and red boxes regions where it is significant after false discovery rate correction for multiple comparisons.

**Supplementary figure 4.**
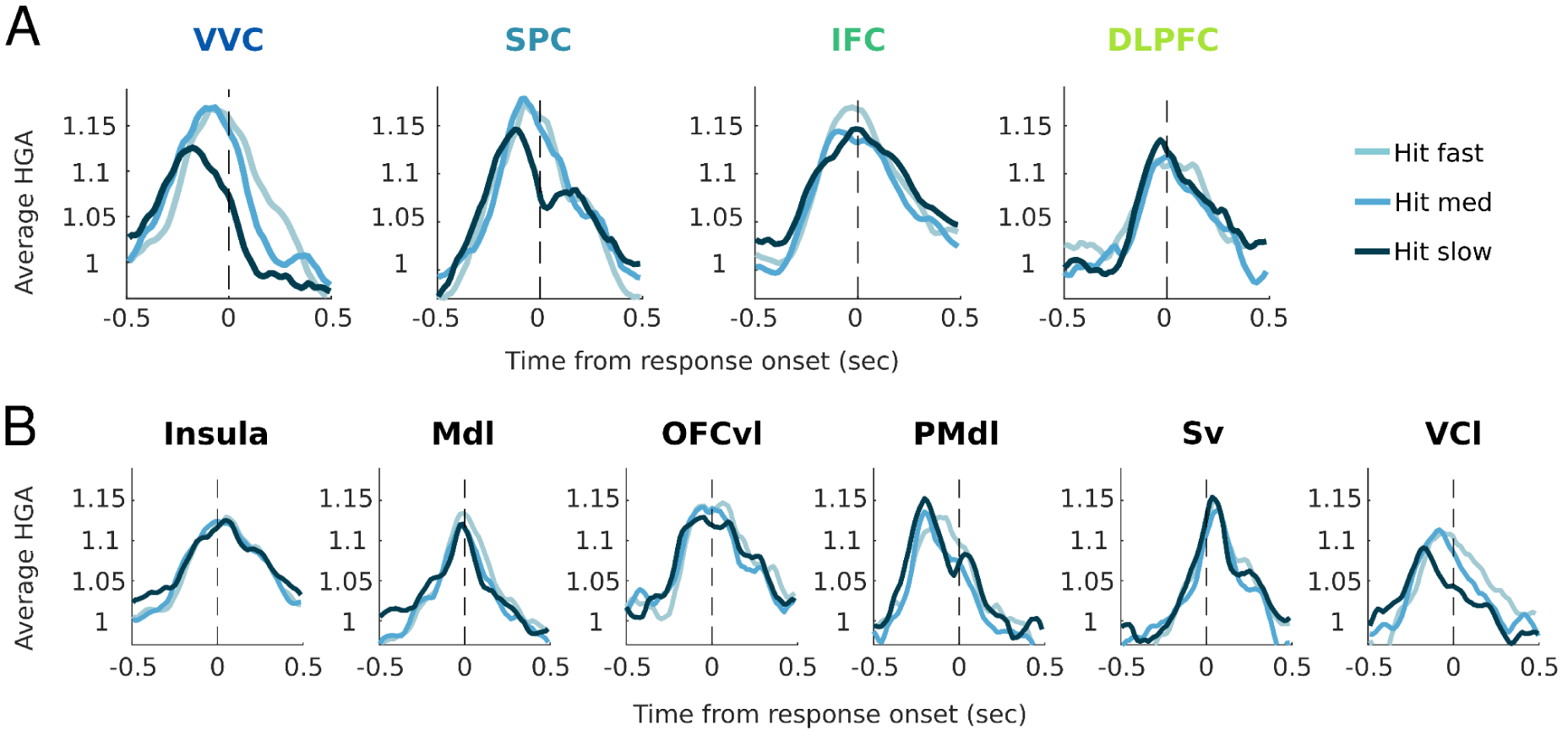
*A.* The averaged activity locked on the response across hit trials separated in three terciles based on reaction times in the four ROIs. B. Same plot for other brain areas that showed a significant effect in *Supplementary figure 3*.

**Supplementary figure 5.**
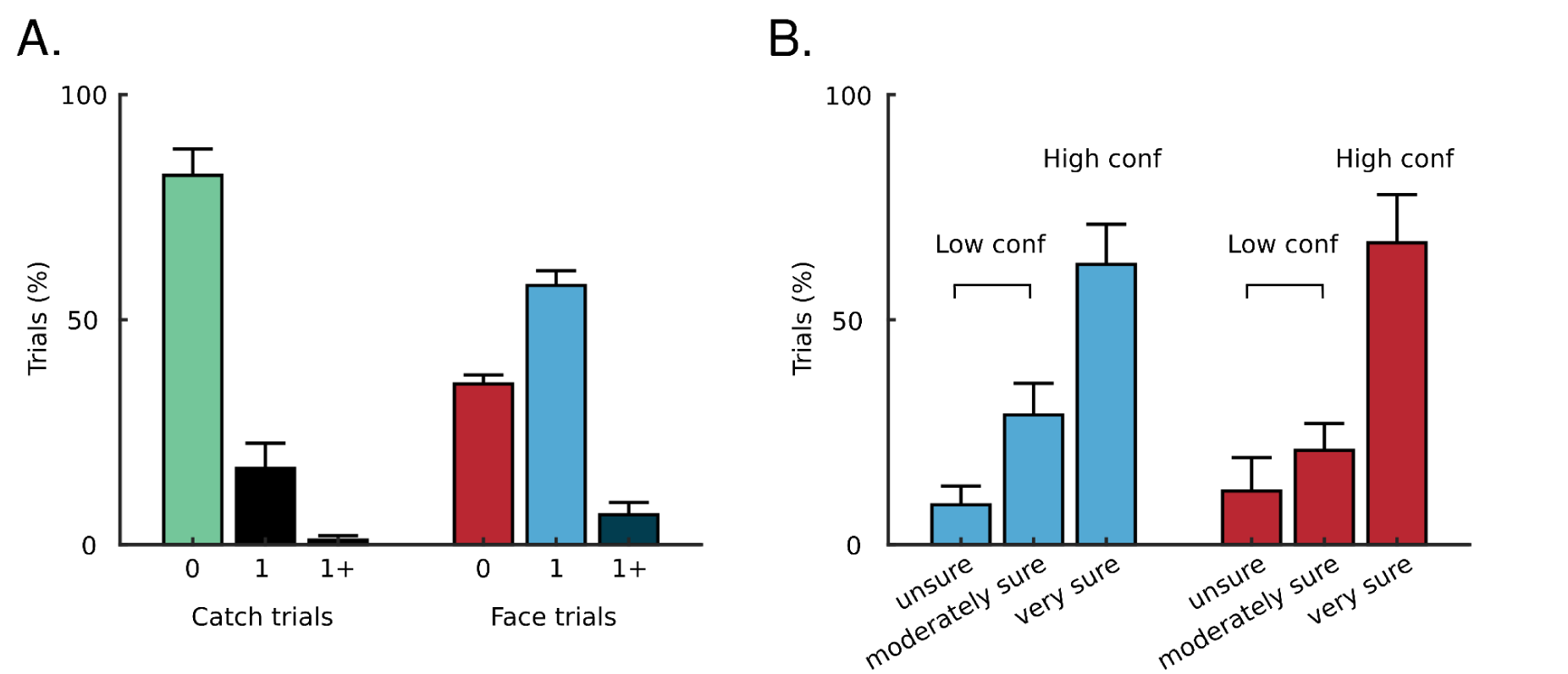
*A.* Average detection reports in experiment 2 for trials in which no face was presented (catch trials) and trials in which one face was presented (face trials). Participants were given the option to report that more than one face was presented (1+ responses). *B.* Subsequent confidence judgments in hit (blue) and miss (red) trials. Participants responded that they were “unsure”, “moderately sure” or “very sure” of their detection response. The first two judgments were pooled as “low confidence” judgments in further analyses.

**Supplementary figure 6.**
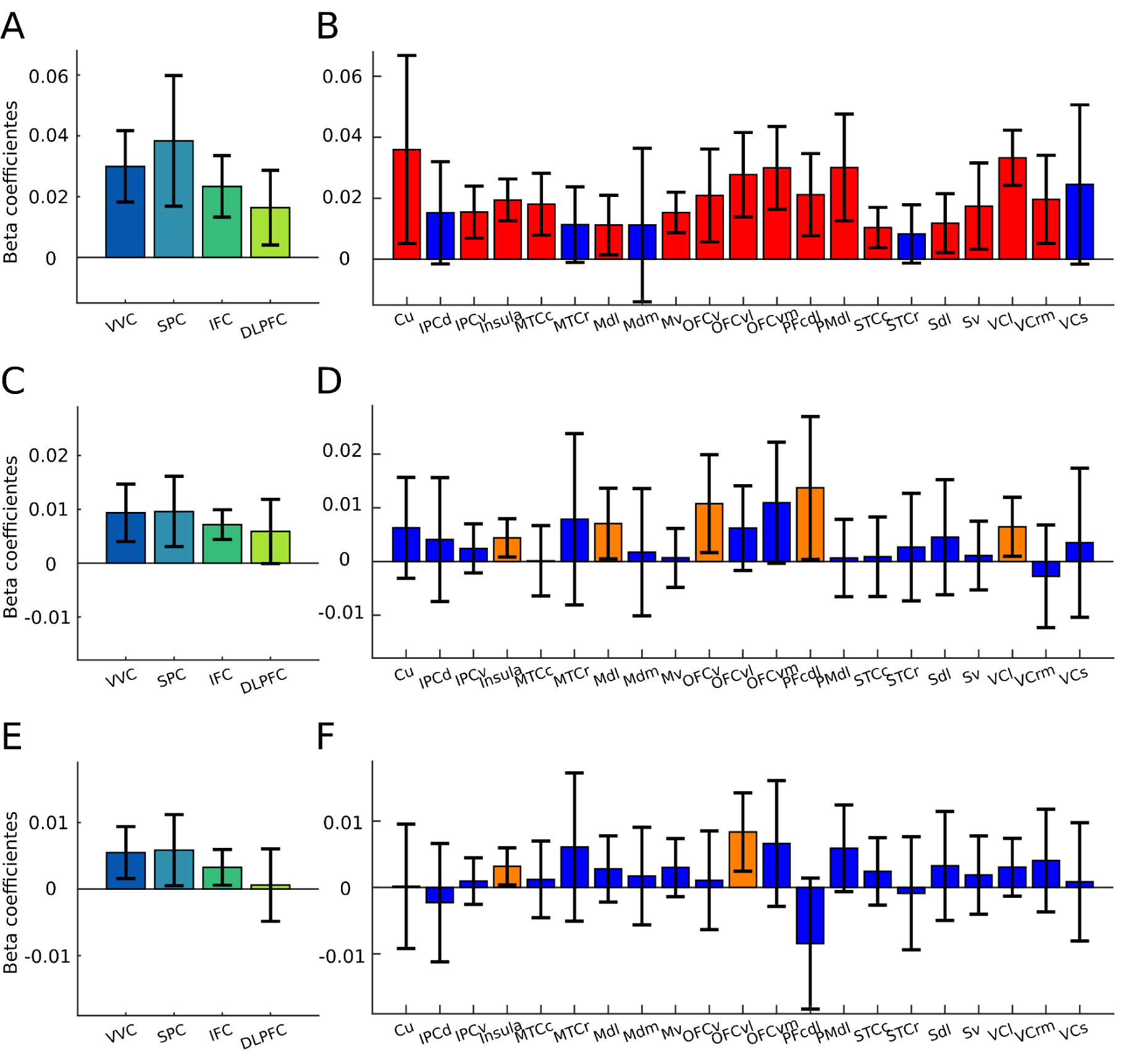
*A.* Beta coefficients for the main effect of detection responses in the analysis of HGA as a function of detection responses and stimulus intensity in the four ROIs in experiment 2. Error bars represent 95% intervals. *B.* Same plot as A for other brain areas. Blue boxes indicate regions where the effect of detection is non-significant, orange boxes regions where it is significant (uncorrected) and red boxes regions where it is significant after false discovery rate correction for multiple comparisons. *C* and *D*: Same plots for the coefficients of the main effect of intensity (threshold vs. suprathreshold). *E* and *F*: Same plots for the coefficients of the interaction between response and intensity.

**Supplementary figure 7.**
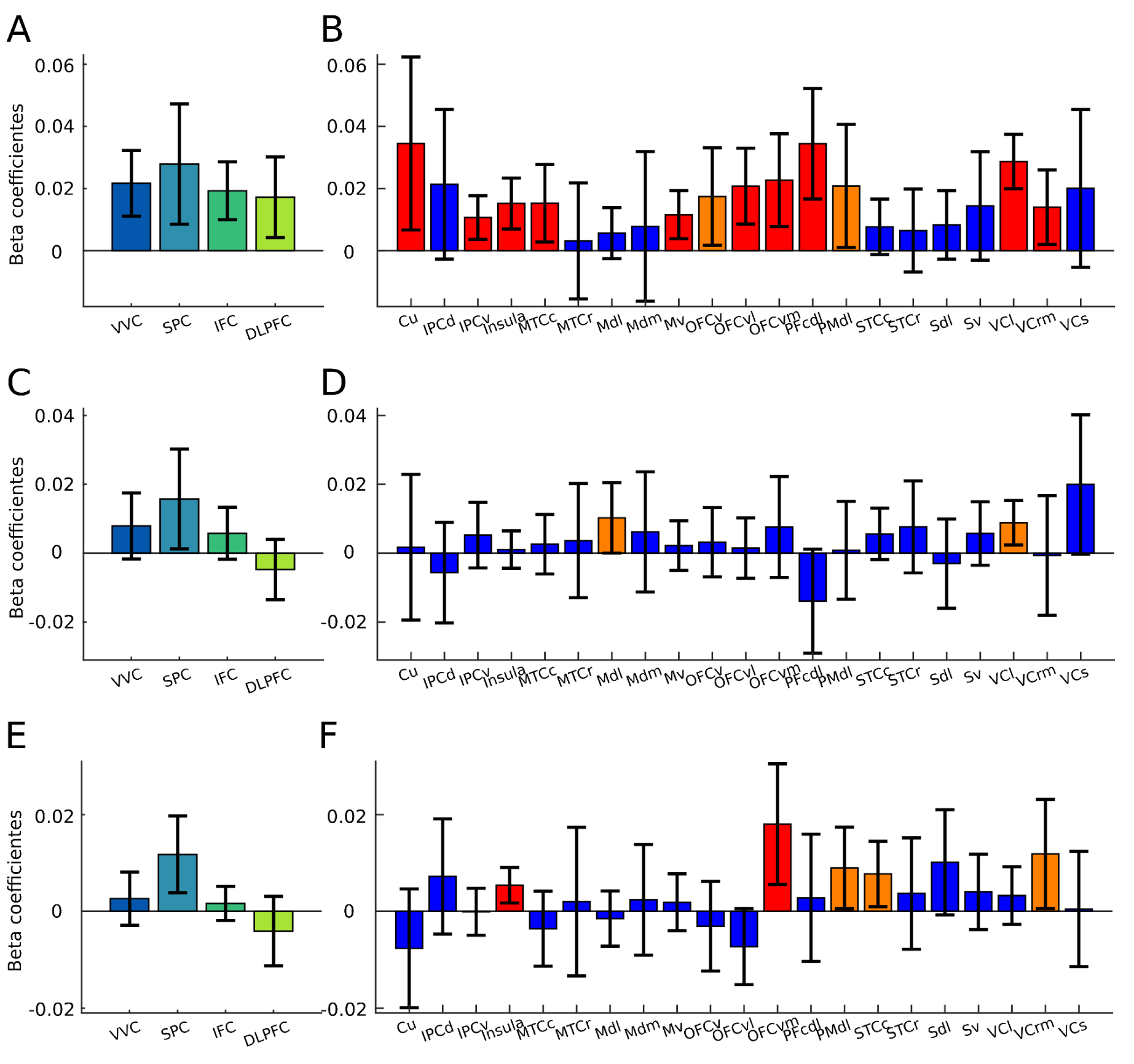
*A.* Beta coefficients for the main effect of detection responses in an analysis of HGA as a function of detection responses and absolute intensity within threshold trials in the four ROIs in experiment 2. The intensity was adapted using a staircase procedure during the experiment (see *Methods*), leading to slightly different absolute values across trials. This analysis shows that the effect of detection remained when absolute intensity was accounted for. Error bars represent 95% intervals. *B.* Same plot as A for other brain areas. Blue boxes indicate regions where the effect of detection is non-significant, orange boxes regions where it is significant (uncorrected) and red boxes regions where it is significant after false discovery rate correction for multiple comparisons. *C, D.* Same plots as A and B, but for the effect of z-scored absolute intensity in the same model. *E, F.* Same plot for the interaction between detection and absolute intensity.

**Supplementary figure 8.**
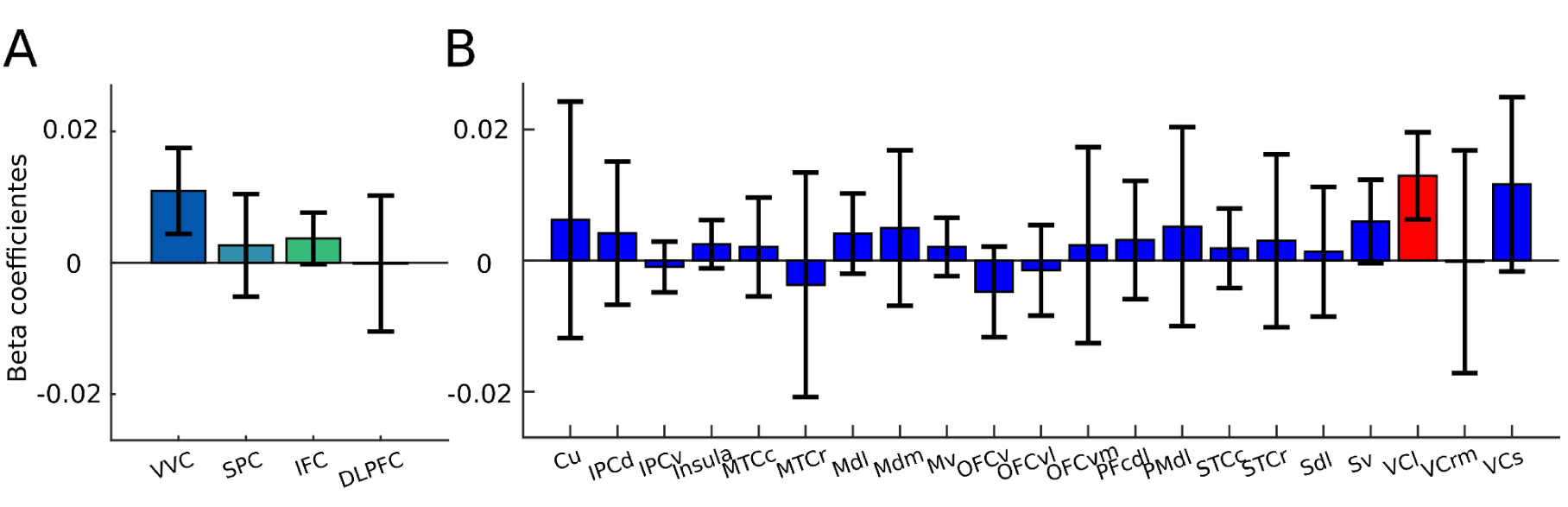
*A.* Beta coefficients for the analysis of HGA as a function of stimulus intensity in the four ROIs in experiment 3. Error bars represent 95% intervals. *B.* Same plot as A for other brain areas. Blue boxes indicate regions where the effect of intensity is non-significant and red boxes regions where it is significant after false discovery rate correction for multiple comparisons.

**Supplementary figure 9.**
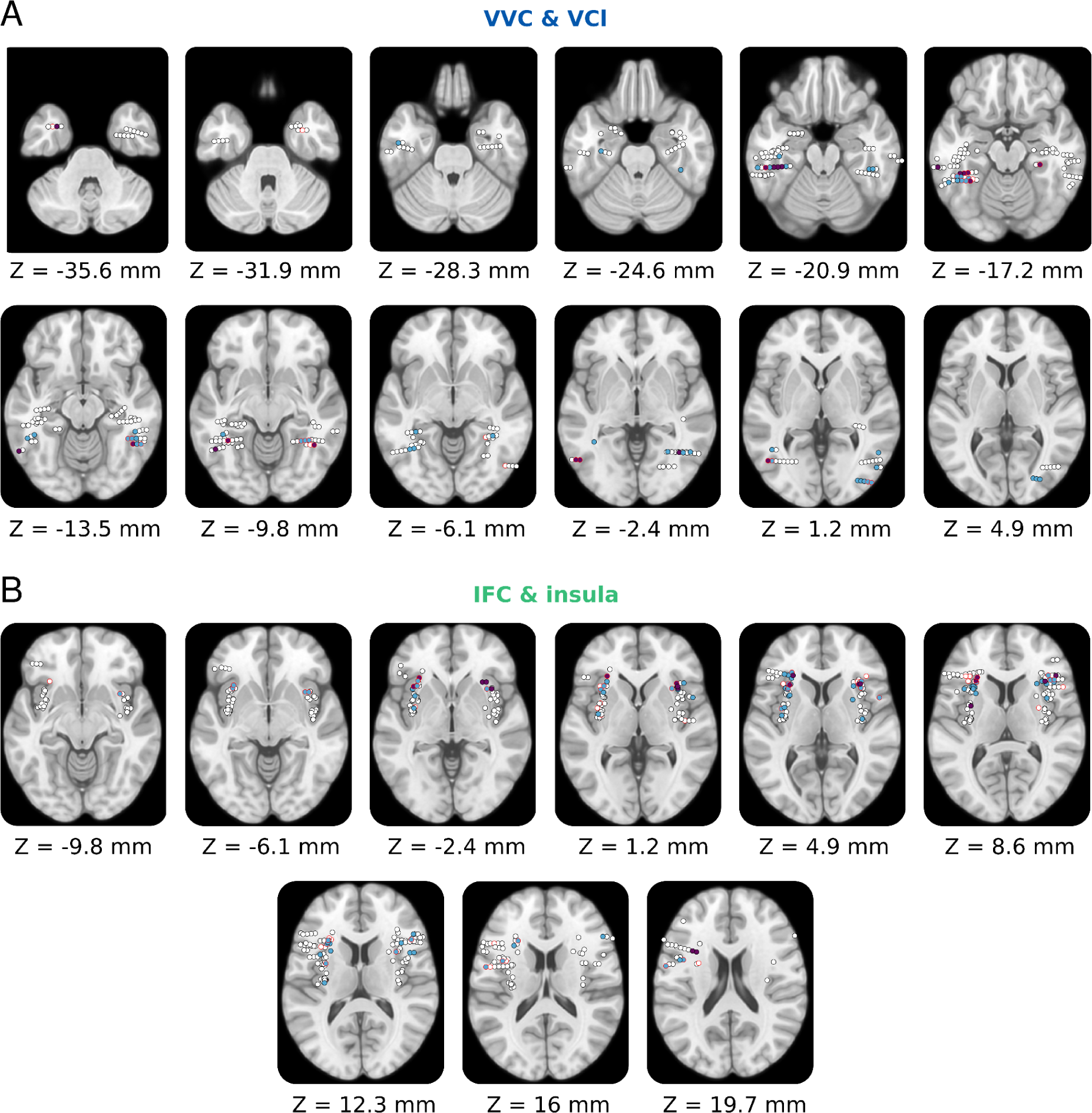
*A.* Projections of channels localized in the VVC and lateral visual cortex (VCl) on transverse planes of the MNI template brain, following the same conventions as on **Figure 4**. *B.* Same for the IFC and the insula.

**Supplementary figure 10.**
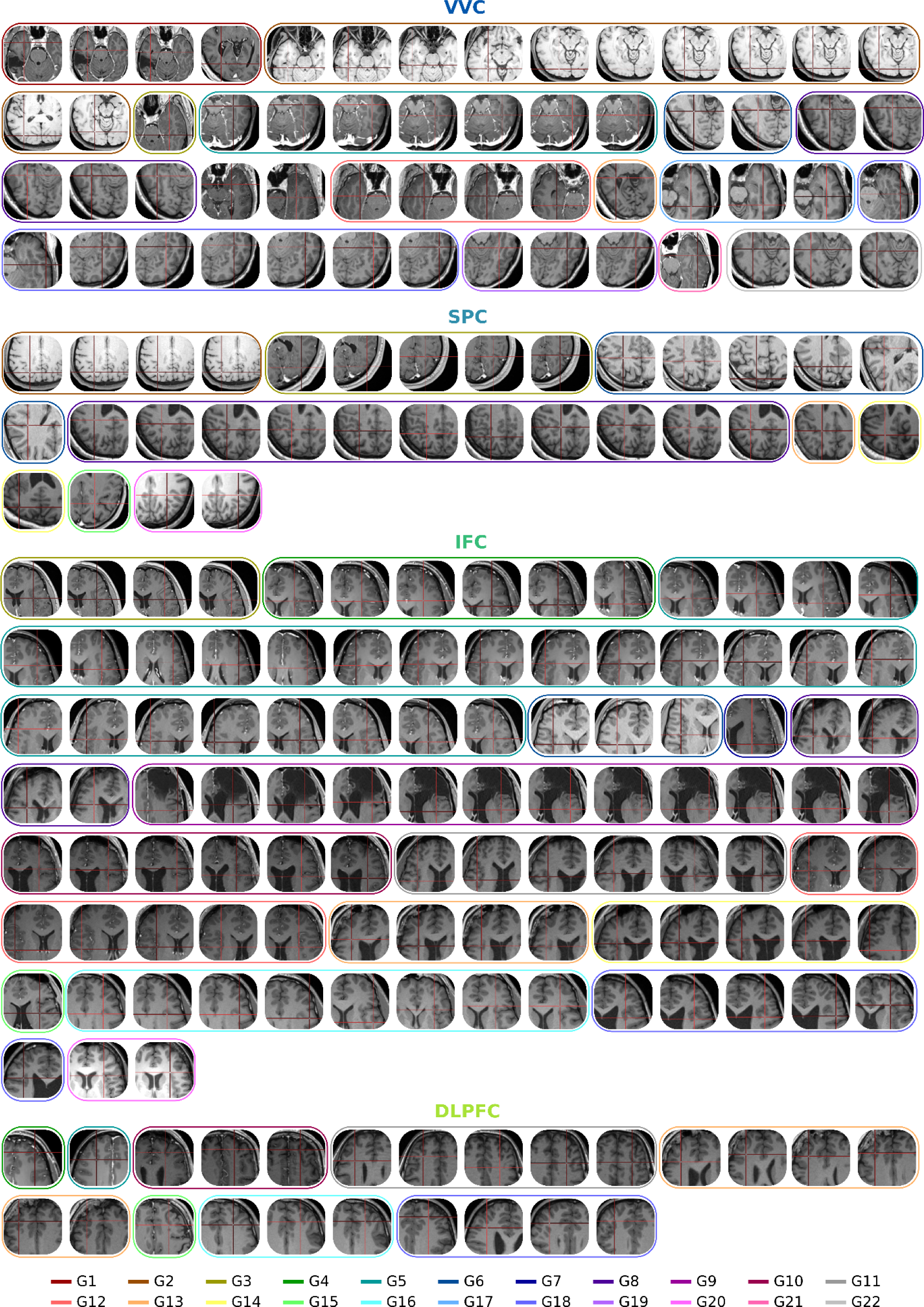
Native localizations of responsive channels for G1-G22 on a transverse plane.

**Supplementary figure 11.**
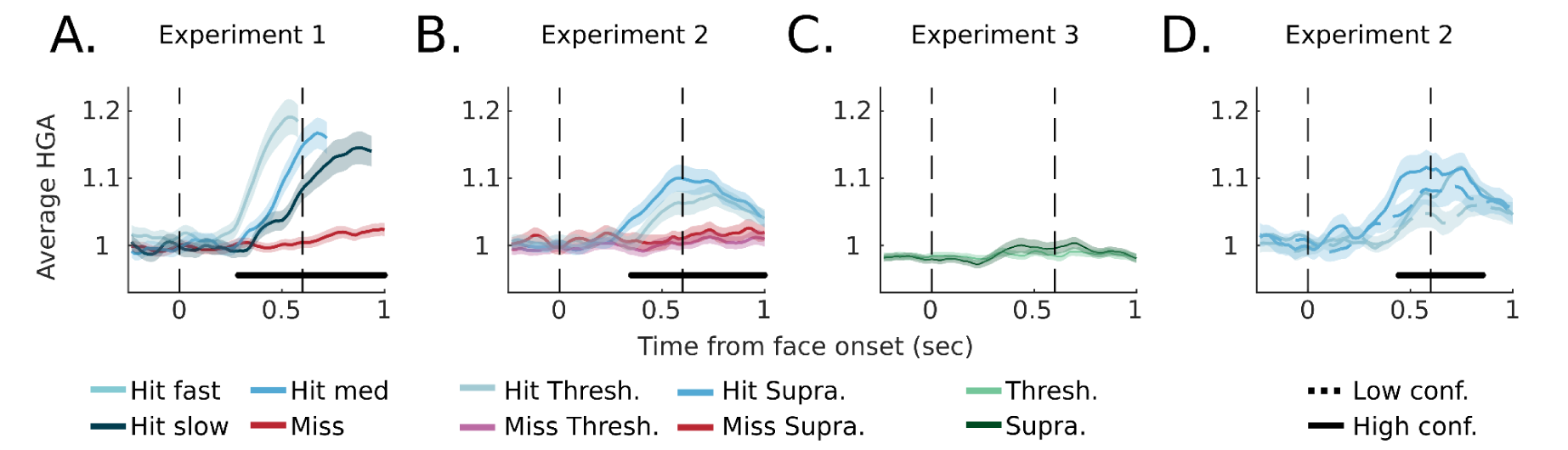
Time-resolved results in a control region encompassing channels in the anterior insular cortex and the internal operculum. Same legend as in *supplementary figure 3*.

**Supplementary figure 12.**
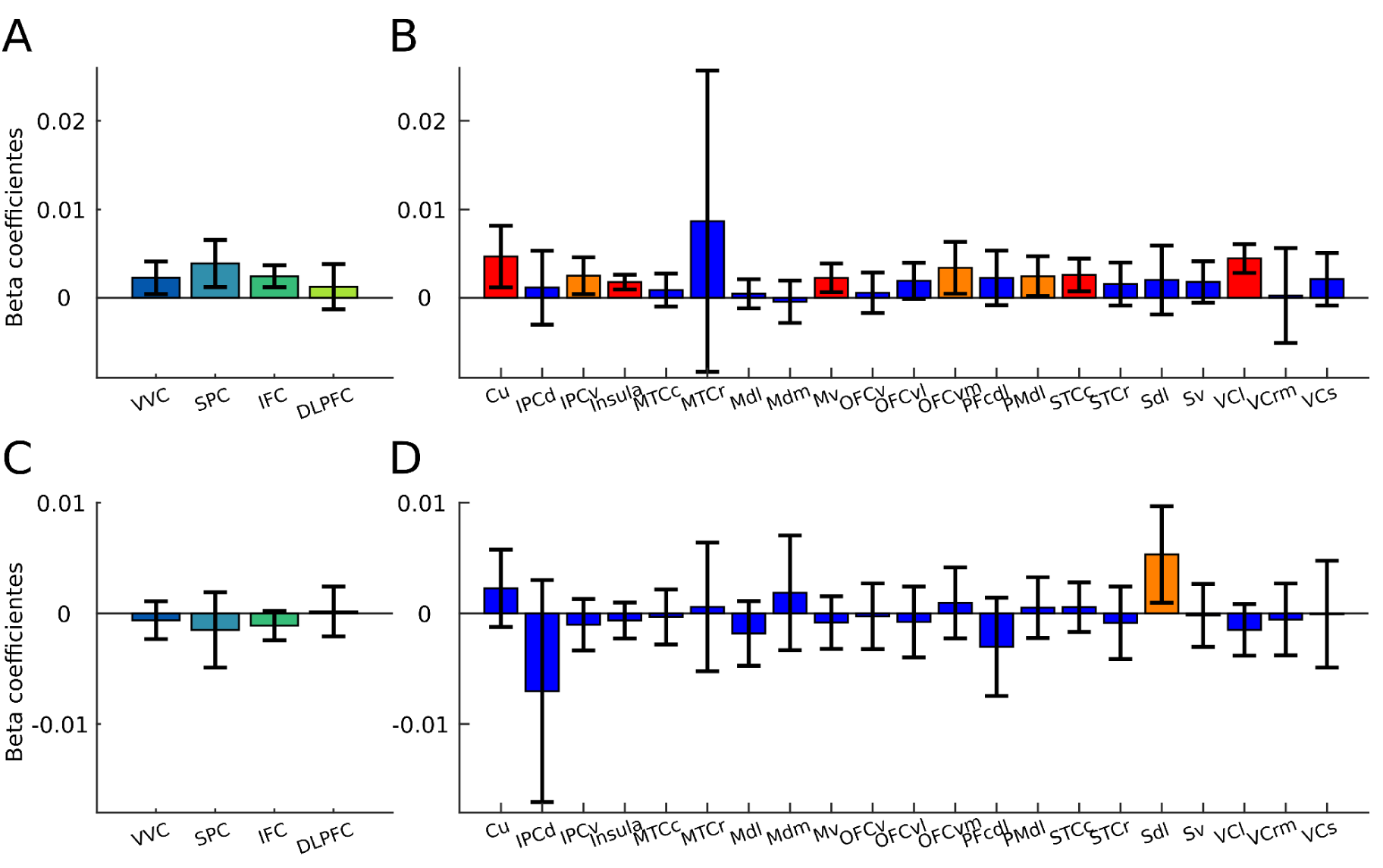
*A.* Beta coefficients for the main effect of confidence (high vs. low) in the analysis of HGA as a function of confidence and stimulus intensity in hit trials in the four ROIs in experiment 2. Error bars represent 95% intervals. *B.* Same plot as A for other brain areas. Blue boxes indicate regions where the effect of confidence is non-significant, orange boxes regions where it is significant (uncorrected) and red boxes regions where it is significant after false discovery rate correction for multiple comparison. *C, D.* Same plots as A and B, but for the effect of confidence in the same model on miss trials.

**Supplementary figure 13.**
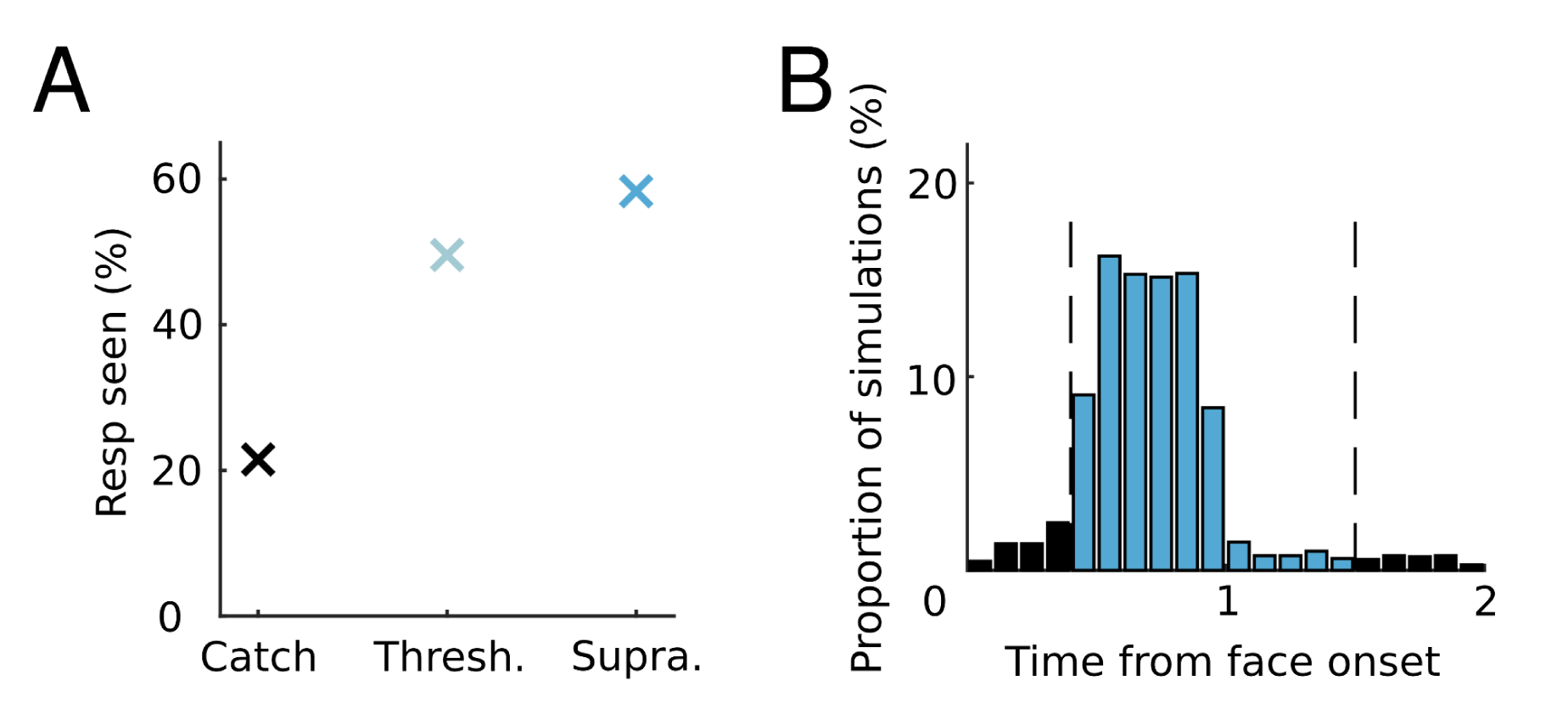
Model simulations. *A.* The results reproduced by the model for detection rates at the three possible intensities. *B.* Reaction times reproduced by the model in the threshold condition.

